# Genetic architecture underlying variation in floral meristem termination in *Aquilegia*

**DOI:** 10.1101/2021.10.18.464884

**Authors:** Ya Min, Evangeline S. Ballerini, Molly B. Edwards, Scott A. Hodges, Elena M. Kramer

**Author notes:** Department of Ecology and Evolutionary Biology, University of Connecticut, Storrs, CT, USA.

## Abstract

Floral organs are produced by floral meristems (FMs), which harbor stem cells in their centers. Since each flower only has a finite number of organs, the stem cell activity of a FM will always terminate at a specific time point, a process termed floral meristem termination (FMT). Variation in the timing of FMT can give rise to floral morphological diversity, but how this process is fine-tuned at a developmental and evolutionary level is poorly understood. Flowers from the genus *Aquilegia* share identical floral organ arrangement except for stamen whorl numbers (SWN), making *Aquilegia* a well-suited system for investigation of this process: differences in SWN between species represent differences in the timing of FMT. By crossing *A. canadensis* and *A. brevistyla*, quantitative trait locus (QTL) mapping has revealed a complex genetic architecture with seven QTL. We identified potential candidate genes under each QTL and characterized novel expression patterns of select candidate genes using *in situ* hybridization. To our knowledge, this is the first attempt to dissect the genetic basis of how natural variation in the timing of FMT is regulated and our results provide insight into how floral morphological diversity can be generated at the meristematic level.

## INTRODUCTION

Indeterminate growth is the foundation of development in all vascular plants and is achieved by the persistent activity of stem cells in the meristems (Steeves & Sussex, 1989). Apical meristems in the shoots and roots are highly organized structures that maintain a delicate, yet robust, balance between the production of cells that give rise to organs and the renewal of the stem cell population. In the flowering plants, when a plant enters the reproductive phase, the vegetative meristem transitions from producing leaves to floral organs. Although the overall cellular organization of the vegetative meristem and the floral meristem (FM) is highly similar, the transition to the FM identity is accompanied by a number of changes in the properties of the meristem, including changes in the rate of organ production, the patterns of primordia initiation, and an eventual loss of indeterminacy.

The loss of indeterminacy (or equivalently, the establishment of determinacy) in the FM is a well-regulated process, termed floral meristem termination (FMT), which is crucial and universal to the development of all flowers (Fig. 1). A typical flower has four types of floral organs: sepals, petals, stamens, and carpels, which are arranged from the outermost to the innermost positions of a flower (Fig. 1). Although the central stem cells stay active in the initial phase of floral organ primordia initiation, this activity will cease at a specific time point, after which all cells will be incorporated into the development of the inner most organs, the carpels (Steeves & Sussex, 1989). The precise control of FMT is critical to ensuring that the flower has the correct number of organs, and variation in the timing of FMT is an important source of floral morphological diversity and novelty. For some species, such as *Arabidopsis thaliana*, FMT occurs relatively quickly, and only four whorls of floral organs are produced. In many other taxa, FM activity is maintained for a more extended period; species from the Magnoliaceae, Monimiaceae, Nelumbonaceae, Nymphaeaceae, Papaveraceae, and Ranunculaceae, for instance, can have hundreds of spirally-arranged or whorled floral organs (Endress, 1990; Fig. 1). Moreover, increased numbers of floral organs can create the raw materials for the evolution of new organ types, such as the sterile staminodes observed in *Aquilegia* (Ranunculaceae) or *Mentzelia* (Loasaceae) (Walker-Larsen & Harder, 2000). Understanding how FMT is regulated in different angiosperm lineages is, therefore, interesting from both developmental and evolutionary perspectives. The diversity in floral morphology of the ~400,000 angiosperm species is seemingly infinite, but one of the few major evolutionary trends independently observed in the transition from early-diverging angiosperms to either the core eudicot or monocot lineages is the transition from variable to stable whorl numbers in a flower (Endress, 1990), which is directly determined by the timing of FMT.

**Figure 1:**
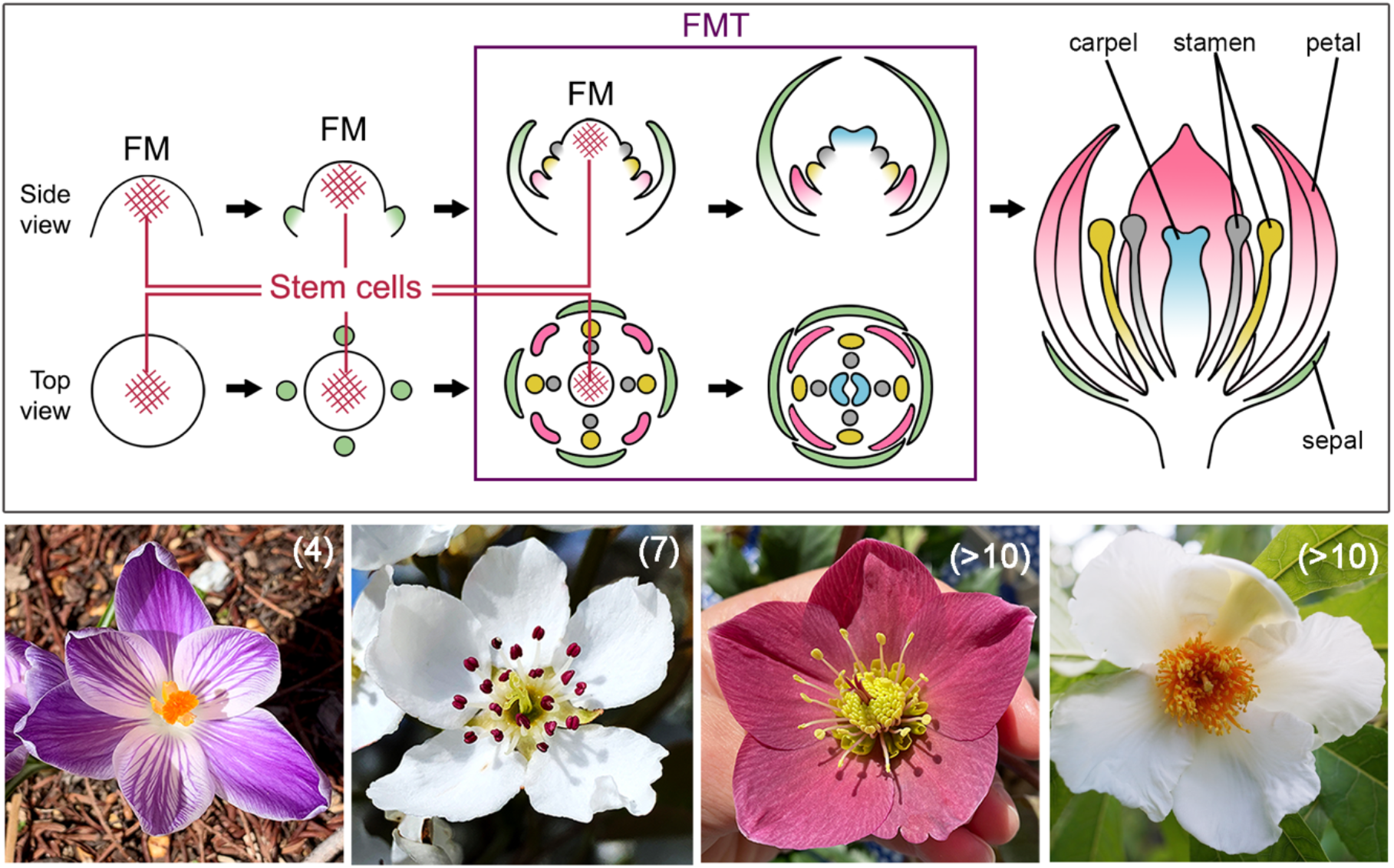
FMT is an important and fine-tuned developmental process that occurs in all flowers. Upper panel: diagram of floral organ initiation and FMT during flower development. Organs of the same whorl share the same colors. Lower panel: example of four flowers with different whorl numbers. From left to right: *Crocus vernus* ‘Pickwick’, *Pyrus communis*, *Helleborus orientalis*, and *Franklinia alatamaha*. Numbers in the parentheses indicate the number of whorls of floral organs in each flower. Photos of *Crocus* and *Pyrus* were taken by Ya Min, and photos of *Helleborus* and *Franklinia* were taken by Evangeline S. Ballerini and Jacob Suissa, respectively

Currently, we have relatively good knowledge of the genes that are responsible for maintaining and terminating stem cell activities in the *A. thaliana* FM, but how the timing of FMT is fine-tuned at the developmental and evolutionary level is poorly understood. In the FM, the maintenance of the stem cell population is achieved by a feedback loop between the homeodomain protein WUSCHEL (WUS) and the CLAVATA (CLV) ligand-receptor system, in which WUS promotes central stem cell activity and induces the expression of the CLV3 peptide, while activation of the CLV signaling pathway, in turn, represses the expression of *WUS* (Schoof *et al.*, 2000; Lenhard, 2003; Müller *et al.*, 2006). In the early developing FM, the expression of the C class organ identity gene *AGAMOUS* (*AG*) is induced by WUS acting as a co-factor with the FM identity protein LEAFY. *AG*, in turn, specifies the identity of stamens and carpels and is also responsible for the down-regulation of *WUS* expression (Lenhard *et al.*, 2001). While the broad conservation of the *WUS*-*CLV* and *WUS*-*AG* feedback loops have been demonstrated in diverse plant taxa (Nardmann & Werr, 2006; Litt & Kramer, 2010; Whitewoods *et al.*, 2020), the exact mechanisms by which AG controls the precise timing of *WUS* down-regulation have only been investigated in *A. thaliana* and tomato (*Solanum lycopersicum*). Specifically, AG activates the expression of a C2H2 zinc-finger transcription factor, *KNUCKLES* (*KNU*), which directly represses *WUS* together with adaptor proteins from the MINI ZINC FINGER (MIF) protein family (Payne, 2004; Sun *et al.*, 2009; Bollier *et al.*, 2018). Accurate timing of FMT is achieved in *A. thaliana* because the activation of *KNU* by AG takes approximately two rounds of cell divisions, during which the stamen primordia are initiated. After FMT is achieved, all of the cells remaining in the center of the FM are incorporated into the carpel primordia (Sun *et al.*, 2009).

However, integral to the precise mechanism of *KNU* activation is that it only allows for the production of one whorl of stamens and one whorl of carpels in a flower. This begs the question of whether this pathway is conserved in systems that have more than four whorls of floral organs. Nonetheless, all currently established model systems (and their close relatives) belong to lineages in the core eudicots or monocot grasses that exhibit no variation in their floral whorl numbers, while most of the plant taxa exhibiting variation in whorls do not have the genomic or genetic resources nor functional tools. Thus, they cannot provide a useful starting point for the investigation of the degree of conservation in this known pathway, or how natural variation in the timing of FMT is regulated in general.

To that end, species in the genus *Aquilegia*, a member of the buttercup family (Ranunculaceae), are well-suited for investigating this fundamental developmental process. There are approximately 70 *Aquilegia* species, which share low interspecific sequence divergence and a high degree of interfertility due to having arisen via a recent adaptive radiation (Filiault *et al.*, 2018). In addition, a number of genetic and genomic resources have been developed in in *Aquilegia* (Kramer, 2009; Filiault *et al.*, 2018). All *Aquilegia* species (with the exception of *A. jonesii*, which lacks staminodes; Munz, 1946) have the same floral organ arrangement, consisting of one whorl of sepals, one whorl of petals, many whorls of stamens, two whorls of staminodes (i.e., 10 in total), and one whorl of carpels (Fig. 2a). All floral organs are produced in a whorl of five, and they are arranged in 10 orthostichies (vertical rows of organs), with alternate orthostichies either positioned directly above the sepals or the petals (Fig. 2a). Therefore, while the number of whorls of sepals, petals, staminodes (except for *A. jonesii*), and carpels is consistent across the genus, the number of stamen whorls varies both within and between species.

**Figures 2:**
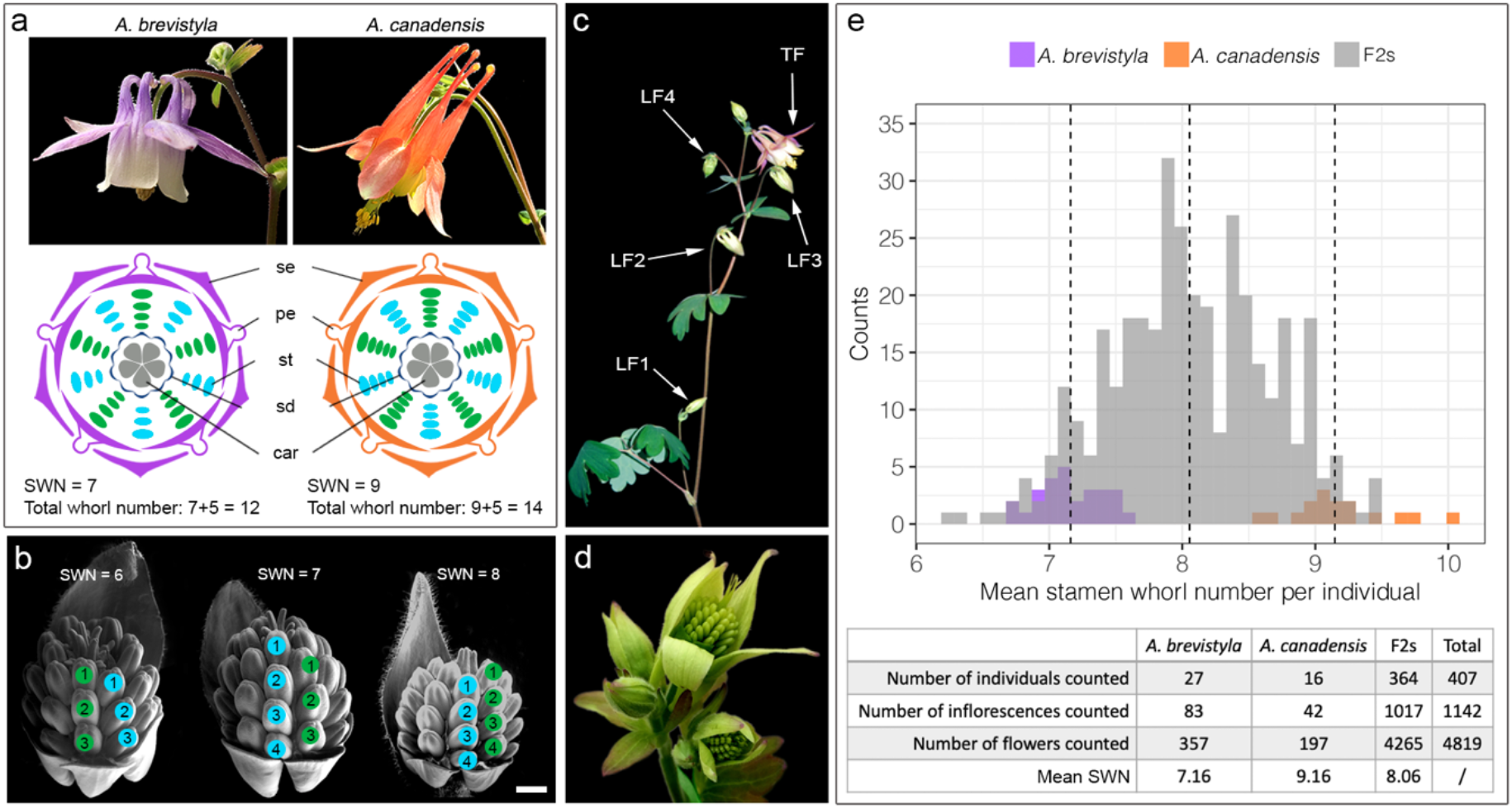
Phenotyping SWN in the parental and F2s populations. (a) Photos of flowers and floral diagrams of *A. brevistyla* and *A. canadensis*. (b) Examples of three F2 flower buds with different SWN. (c) Flowers that were sampled per inflorescence. (d) Developmental stages for which SWN were counted. (e) Histogram and summary statistics of SWN distribution in parental species and the F2s. In (a) and (b), stamen whorls positioned above the sepals are colored in blue while stamen whorls positioned above the petals are colored in green. Scale bar in (b) = 100 μm. se: sepals; pe: petals: st: stamens; sd: staminodes; car: carpels

*Aquilegia brevistyla* and *A. canadensis* are North American sister species (Bastida *et al.*, 2010; Fior *et al.*, 2013; Fig. 2a). Given that the number of whorls of all the non-stamen floral organs is identical in these two species, the variation in what we term Stamen Whorl Number (SWN) can be directly translated into variation in the timing of FMT: if the FM is terminated earlier, it will have a smaller SWN compared to a flower of which the FM stays active longer. Using SWN as a quantitative trait, we crossed *A. brevistyla* and *A. canadensis* and conducted quantitative trait locus (QTL) mapping in the resultant F2 generation. Our results reveal a complex genetic architecture with seven major QTL underlying the variation of FMT. To our knowledge, this is the first study to dissect the genetic basis of how natural variation in the timing of FMT is regulated, and our results highlight several potential candidate genes and molecular pathways that may contribute to the regulation of FMT in *Aquilegia*.

## RESULTS

### SWN variation in the parental species and the F2s

We counted the SWN from 357, 197, and 4265 flowers from 27, 16, and 364 *A. brevistyla*, *A. canadensis*, and F2 individuals, respectively (Fig. 2). The SWN per individual of the parental species did not overlap: the mean SWN of *A. brevistyla* ranges from 6.69 to 7.57; that of *A. canadensis*, from 8.54 to 10; and that of the F2s, which overlapped with the range of both parental species, from 6.20 to 9.50 (Fig. 2e). The mean SWN for all *A. brevistyla*, *A. canadensis*, and F2s were 7.16, 9.16, and 8.06, respectively. Subsequently, we analyzed whether the position of flowers on the inflorescence is associated with their SWN (Fig. 2c). Flower position had a significant effect on the SWN for both parental species but not for the F2s (Fig. S1). Given lower germination rates of interspecific hybrid seeds, seeds generated by selfing five different F1 plants (all of which had the same parents) were used to get 364 F2 progeny that reached the flowering stage (n=53-81 per F1). One-way ANOVA revealed that the SWN of the F2s differed significantly between the F1 parents (Fig. S2). Lastly, we also analyzed the variation of SWN among flowers of the same plants. Interestingly, a small portion of *A. brevistyla* (7.4%), *A. canadensis* (18.8%), and F2s (6%) showed no variance in the SWN across all flowers counted within an individual, and this phenomenon was dependent on neither the number of flowers counted per individual plant (Fig. S3; Pearson’s correlation = 0.052, t = 0.98418, df = 345, *p* = 0.3257) nor the F1-parent-of-origin of the F2s (Fig. S3). No significant correlation between the mean SWN per individual and the standard deviation of SWN per individual was detected (Pearson’s correlation = 0.035, t = 0.70438, df = 404, *p* = 0.4816).

### Floral meristem size

To determine whether the initial FM sizes were different between the parental species, we measured the widths of FMs of the parental species at their earliest developmental stages (Fig. 3). In general, the FMs of *A. canadensis* appeared to be slightly, but significantly, wider than those of *A. brevistyla* throughout the early developmental stages (Fig. 3; Table S1). The average FM widths of *A. brevistyla* and *A. canadensis* before the initiation of carpel primordia were μm and 191.67 μm, respectively (Table S1). Interestingly, the temporal developmental windows for significant FM size expansion seemed to be longer in *A. canadensis* than *A. brevistyla* (Fig. 3; Table S1). The significant increase in widths of *A. brevistyla* FMs occurred when there were 0 to 4 whorls of non-sepal floral organs initiating, while the significant increase in the widths of *A. canadensis* FMs encompassed a larger developmental period, ranging from the stages that there were 0 to 6 whorls of non-sepal floral organs initiating (Table S1).

**Figure 3.**
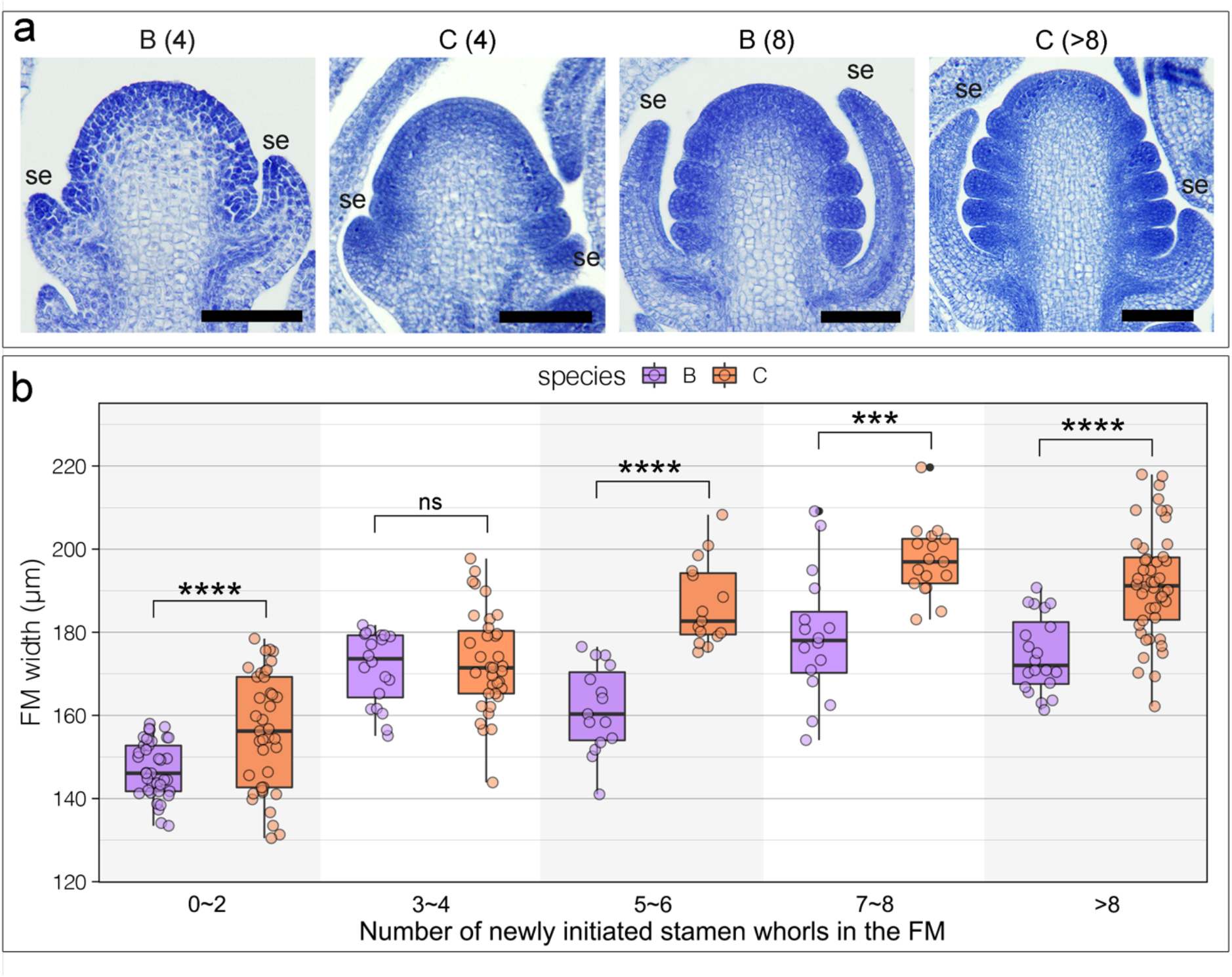
FM widths measurements of the parental species during the early developmental stages. (a) Examples of FM morphologies in *A. brevistyla* and *A. canadensis* at early developmental stages. Numbers in the parenthesis indicate the number newly initiated stamen whorls in each FM. Scale bars = 100 μm. (b) Comparison of FM width of difference developmental stages between *A. brevistyla* and *A. canadensis*. Each data point represents a measurement of a FM width from a section. Three to six continuous sections were measured for each FM, and at least three FMs were measured for each developmental stage of each species. Comparison of FM widths of each stage between the parental species was done using Tukey’s HSD. ns: not significant; ***: p-value < 0.001; ****: p-value < 0.0001; se: sepal; B: *A. brevistyla*; C: *A. canadensis*.

### Genetic architecture underlying stamen whorl variation

The genetic map was constructed using a total of 620 genetic markers, which fell into seven linkage groups, matching the n=7 chromosomes in the *Aquilegia* genome (Fig. S4). We recovered seven major QTL using the mean SWN per individual as a phenotype and the F1-parent-of-origin as a covariate, with one QTL on each chromosome (Fig. 4a; Table 1). The difference in LOD scores between models that included or excluded the covariate are diminutive on all chromosomes (ranging from −0.3 to 0.49; Fig. S5a), indicating no significant interaction between the QTL and the covariate. While several LOD profiles from a single QTL model suggested that there may be two QTL on a chromosome, a two-dimensional genome scan failed to detect evidence for the presence of more than one QTL on a single chromosome (Table S2). The presence of only one true QTL on chromosome 2 was further confirmed by controlling the two potential QTL: when the true QTL (i.e., Q2) was controlled, the presence of the second peak also disappeared (Fig. S5b). We did detect significant interactions between two pairs of QTL: Q3 and Q7, and Q1 and Q6 (Fig. S5c), and thus incorporated these interactions in the full QTL model (Table 1).

**Table 1:** Summary statistics for QTL. Chr: chromosome; PVE: percent variance explained; Add./(a): additive affects; Dom./(d): dominant effects.

| Full model result: |  |  |  |  |  |  |
| --- | --- | --- | --- | --- | --- | --- |
| $y \sim Q1 + Q2 + Q3 + Q4 + Q5 + Q6 + Q7 + Q1:Q6 + Q3:Q7$ | | | | | | |
|  | df | SS | MS | LOD | PVE | Pvalue(F) |
| Model | 22 | 61.87 | 2.81 | 48.04 | 44.47 | 0 |
| Error | 331 | 71.27 | 0.22 |  |  |  |
| Total | 353 | 133.14 |  |  |  |  |

**Table 1: Summary statistics for QTL.** Chr: chromosome; PVE: percent variance explained;
| Drop one QTL at a time ANOVA table |  |  |  |  |  |  |  |  |  |  |
| --- | --- | --- | --- | --- | --- | --- | --- | --- | --- | --- |
|  | Chr | Position (cM) | df | Type III SS | LOD | PVE | F value | Pvalue (F) | Add. | Dom. |
| Q1 | 1 | 44.233 | 6 | 5.87 | 6.1 | 4.4 | 4.6 | <0.001 | 0.05 | -0.16 |
| Q2 | 2 | 19.676 | 2 | 4.89 | 5.1 | 3.7 | 11.4 | <0.001 | -0.07 | 0.2 |
| Q3 | 3 | 24.050 | 6 | 7.43 | 7.6 | 5.6 | 5.7 | <0.001 | -0.11 | -0.01 |
| Q4 | 4 | 25.406 | 2 | 11.73 | 11.7 | 8.8 | 27.3 | <0.001 | -0.27 | 0.07 |
| Q5 | 5 | 30.155 | 2 | 6.89 | 7.1 | 5.2 | 16.0 | <0.001 | -0.25 | 0.17 |
| Q6 | 6 | 34.828 | 6 | 11.72 | 11.7 | 8.8 | 9.1 | <0.001 | -0.14 | -0.27 |
| Q7 | 7 | 20.944 | 6 | 7.38 | 7.6 | 5.6 | 5.7 | <0.001 | 0 | 0.15 |
| Q1 * Q6 |  |  | 4 | 5.45 | 5.7 | 4.1 | 6.3 | <0.001 |  |  |
| Q3 * Q7 |  |  | 4 | 5.39 | 5.6 | 4.1 | 6.3 | <0.001 |  |  |
| Estimated effects for QTL interactions |  |  |  |  |  |  |  |  |  |  |
| Q1 (a) * Q6 (a): 0.01 |  |  | Q1 (d) * Q6 (a): -0.23 |  |  | Q1 (a) * Q6 (d): -0.38 |  |  | Q1 (d) * Q6 (d): 0.45 |  |
| Q3 (a) * Q7 (a): 0.19 |  |  | Q3 (d) * Q7 (a): 0.09 |  |  | Q3 (a) * Q7 (d): -0.16 |  |  | Q3 (d) * Q7 (d): -0.24 |  |

**Figure 4:**
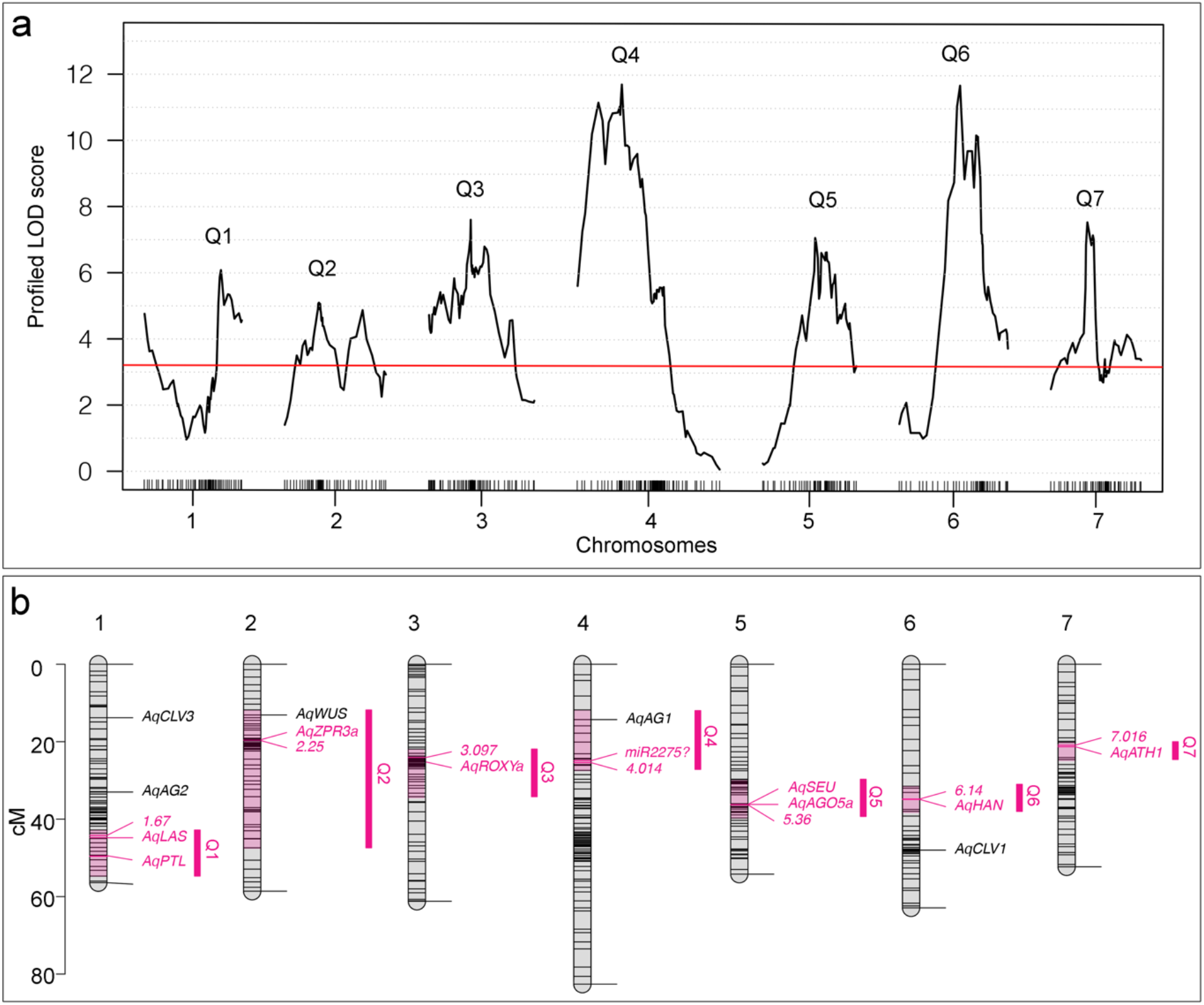
Genetic architecture and candidate genes. (a) LOD scores across seven chromosomes. Red line: α = 0.05 genome-wide significance cutoff based on 1000 permutations. (b) Locations of QTL interval (pink regions on the chromosomes and magenta vertical bars), candidate genes, and genetic markers. All the genetic markers were named in numeric forms (e.g., 1.67 and 2.25) and only markers with the highest LOD scores under each QTL are shown.

The full QTL model had a total LOD score of 48 and explained 46.5% of the observed phenotypic variation (Table 1). Q3, Q4, and Q5 exhibit larger additive effects than dominant effects, while the remaining QTL have larger dominant effects on the phenotypic variation (Fig. 4a, S6; Table 1). The phenotypic variation explained by each QTL and the QTL interactions was very similar, with Q4 and Q6 having the largest additive and dominant effects, respectively, and each explained 8.8% of the phenotypic variation (Table 1). This suggested the genetic architecture of SWN is a complex trait controlled by multiple loci, each with small effect.

### Candidate genes

In order to identify potential candidate genes underlying these QTL, we examined the genomic regions defined by markers that flanked the 95% Bayesian credible interval of each QTL, which was calculated by using the posterior distribution of 10^LOD on a given chromosome. The genomic regions of Q1, Q4, Q6, and Q7 were less than 6 Mb in size, while those of Q2, Q3, and Q5 were more than 20 Mb (Table S3). Among all the QTL, Q6 and Q2 had the smallest (1.5 Mb) and the largest (36.5 Mb) intervals, containing 226 and 3242 genes, respectively (Table S3). We narrowed down the list of candidate genes by using previously published RNA-sequencing (RNAseq) data for early FMs of *A. coerulea* ‘Kiragami’, which sampled developmental stages covering the FMT window (Min & Kramer, 2020). All QTL intervals, with the exception of Q4, had approx. 70% of their total genes expressed in at least one of the RNAseq developmental stages; only 46.21% of the total Q4 genes were expressed in early developmental stages (Table S3).

Subsequently, we sought to identify candidate genes under each QTL based on 1) the locations of the candidate genes relative to the location of the markers with the highest LOD scores, 2) gene expression levels during early FM developmental stages, and 3) homology to previously studied loci related to meristem function (i.e., not just restricted to FM). Because our genotyping method used 0.5 Mb or 1Mb binned genomic regions as the genetic markers rather than single nucleotide markers (see details in Materials and Methods), we gave the highest priority to genes located in the region of the marker with the highest LOD scores.

Within the highest LOD bin of Q1, we identified a homolog of *LATERAL SUPPRESSOR* (*LAS*), *AqLAS*. *LAS* encodes a member of the GRAS family of putative transcriptional regulators, and mutations in the *LAS* orthologs in *A. thaliana* and tomato lead to a loss of axillary meristems (Schumacher *et al.*, 1999; Greb, 2003; Wang *et al.*, 2014). Moreover, one additional gene within the interval that was located 3 Mb away from *AqLAS* also caught our attention: *AqPETAL LOSS* (*AqPTL*). *PTL* is a floral organ boundary gene in *A. thaliana* that controls cell proliferation in a non-cell autonomous fashion (Griffith *et al.*, 1999; Brewer *et al.*, 2004; Lampugnani *et al.*, 2012). We considered *AqPTL* interesting for two reasons. First, PTL has been shown to physically interact with and be transcriptionally regulated by C2H2 transcription factor JAGGED (JAG) (Sauret-Güeto *et al.*, 2013). In addition, silencing of *AqJAG* led to early FM arrest in *A. coerulea*, indicating that it is an important gene in maintaining the *Aquilegia* FM (Min & Kramer, 2017). Second, PTL and the gene product of *HANABA TANARU* (*HAN*) interact by sharing JAG as direct protein partners to regulate floral morphogenesis in *A. thaliana* (Ding *et al.*, 2015), and the homolog of *HAN* is a candidate gene under our Q6 (Fig. 4b).

Interestingly, *AqWUS* is within the 95% Bayesian credible intervals of Q2, located at the edge of the interval (Fig. 4b). Since the expression of *AqWUS* has not been examined *in situ*, we analyzed its expression pattern during the early developmental stages of *A. coerulea* FM (Fig. 5a-e). *AqWUS* is expressed in a small population of cells in the center of the FM from the earliest stages (Fig. 5a-e). The expression in these central zone cells persists during the initiation of floral organs and disappears when carpel primordia start to initiate. These observations are consistent with the expression of *WUS* orthologs in all taxa examined to date (Nardmann & Werr, 2006; Galli & Gallavotti, 2016), suggesting functional conservation of *AqWUS* as well. However, the marginal position of *AqWUS* in Q2 makes it a less compelling candidate.

**Figure 5.**
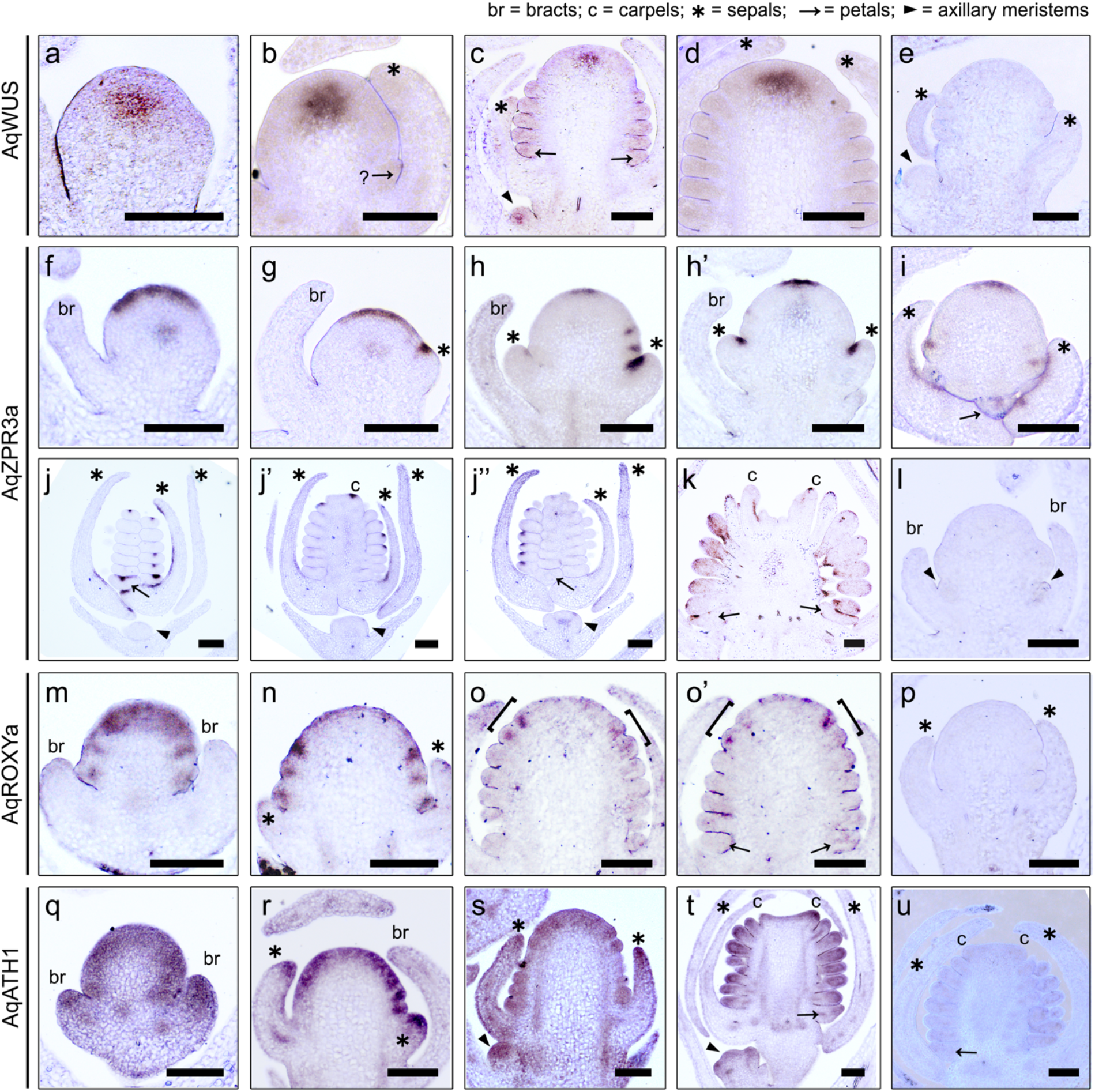
*In situ* hybridization of *AqWUS* and candidate genes. (a-e) Expression patterns of *AqWUS* (a-c) and its negative control (e). (a) A young FM that has not produced any floral organs. (b) A young FM that is in the process of initiating either a petal primordium or the outer most stamen primordia (and thus indicated by an arrow with a question mark). (c) A FM that has at least eight whorls of stamens initiated; *AqWUS* expression can also be seen at the axillary meristem below this FM (arrowhead). (d) A FM that has at least 11 whorls of stamens produced. (e) Sense probe on a FM with 3-4 whorls of stamens. (f-l) Expression patterns of *AqZPR3a* (f-k) and its negative control probe (l). (f) A young FM that has not produced any floral organs. (g) A young FM that has just started to produce sepal primordia (asterisk). (h, h’) Serial sections through the same young FM that has only produced sepal primordia (asterisks). (i) A slightly tangential section through a FM that has produced 1-2 whorls of stamens. (j, j’, j”) Serial sections through the same young floral bud that has initiated all floral organs. Expression of *AqZPR3a* is also e seen in the central zone of the axillary meristem in (j”) (arrowhead). (k) A young floral bud in which all floral organs are differentiating. (l) Meristem at same stage as (f) hybridized with sense probe. (m-p) Expression of *AqROXYa* (m-o’) and its negative control (p). (m) A young FM that has not produced any floral organs. (n) A young FM that is in the process of initiating petal or the outer-most stamen primordia. (o, o’) Serial sections through the same FM showing *AqROXYa* is only expressed on the abaxial side of the newly emerging primordia (brackets). (p) Meristem at same stage as (p) hybridized with sense probe. (q-u) Expression of *AqATH1* (q-t) and its negative control (u). (q) A young FM that has not produced any floral organs. (r) A young FM that is in the process of initiating petal and stamen primordia. (s) A FM initiating stamen primordia and an associated axillary meristem (arrowhead). (t) A floral bud with carpel primordia just initiated and an axillary meristem, which has just initiated the sepal primordia. (u) Meristem at same stage as (t) hybridized with sense probe. All scale bars = 100 μm.

Under Q2, 8.7 Mb away from *AqWUS* within the highest LOD interval, we identified another candidate, the gene *AqLITTLE ZIPPER 3a* (*AqZPR3a*) (Fig. 4). *AqZPR3a* encodes a small-leucine zipper-containing protein and is the homolog of the previously identified gene *ZPR3* in *A. thaliana* (Weits *et al.*, 2019). Homologs of the *ZPR* genes have been shown to regulate leaf polarity and shoot apical meristem maintenance in *A. thaliana* and tomato (Wenkel *et al.*, 2007; Kim *et al.*, 2008; Weits *et al.*, 2019; Xu *et al.*, 2019), but little is known about whether they are involved in any FM-specific functions in these plant systems. In *Aquilegia, AqZPR3a* exhibits fascinating and dynamic expression patterns during early FM development (Fig. 5f-l). At the earliest stages of the FM, before floral organ primordia are initiated, concentrated expression of *AqZPR3a* is detected across the central epidermal layer of the FM, and moderate expression is found in the central zone (Fig. 5f). This strong expression in the FM epidermal layer persists until the FM has initiated several whorls of floral organs, but the width of the domain seems to contract as FM development proceeds (Fig. 5f-i). On the other hand, the expression in the central zone disappears rapidly after initiation of the sepal primordia (Fig. 5f, g, h’, j’’). Strong expression of *AqZPR3a* is also detected at the adaxial boundary of all initiating floral organ primordia (Fig. 5h-j’’). However, these adaxial expression domains are restricted to the median region of the primordia, rather than the entire abaxial surface, which can be seen in serial sections through the same FM in Fig. 5h, h’, as well as in Fig. 5j, j’, j’’. Moreover, strong but patchy expression of *AqZPR3a* is detected in the adaxial epidermal layer of slightly older lateral organs, such as the sepals (Fig. 5i, j, j’), petals (Fig. 5k), stamens (Fig. 5k) and carpels (Fig. 5k). Intriguingly, expression of the *ZRP* genes across the epidermal layer of the early meristems has never been observed in any other plant systems.

Under Q3, we identified the candidate gene *AqROXYa* which codes for a thioredoxin superfamily protein and is a homolog of *A. thaliana* genes *ROXY1* and *ROXY2* (Fig. 4; Fig. S7). In *A. thaliana*, *ROXY1* appears to be a negative regulator of *AG* and functions to regulate petal initiation (Xing *et al.*, 2005; Quon *et al.*, 2017), and *ROXY2* with other members of the thioredoxin family members have been shown to regulate both floral organ development and stress responses (Xing & Zachgo, 2008; Murmu *et al.*, 2010; Li *et al.*, 2019). However, the homolog of *ROXY* in maize has been shown to regulate shoot meristem size and phyllotaxy (Yang *et al.*, 2015). In *Aquilegia*, expression of *AqROXYa* is detected across the FM at the earliest developmental stages (Fig. 5m), but this broad expression disappears once the primordia begin initiating. Likewise, *AqROXYa* is detected in a restricted abaxial region of emerging floral organ primordia but quickly declines once they are initiated (Fig. 5m-o’). For instance, in a FM with several whorls of initiated stamen primordia, the expression of *AqROXYa* is only detected in the abaxial side of the innermost two whorls of emerging organ primordia (Fig. 5o, o’).

The confidence interval of Q4 spanned 3.8 Mb but only contained 176 annotated genes that were expressed in the RNAseq dataset (Table S3, S4). The only loci with homologs that are likely to function in meristems are two closely related tandem duplicates of the *AGAMOUS* homolog *AqAG1* (Fig. 4b). However, *AqAG1* is located at the edge of the genomic interval, quite distant from the marker with the highest LOD score (Fig. 4b). Within the 1Mb region (4,800,000..5,800,000) that contained the marker with the highest LOD, there were only 40 expressed genes, eight of which are *Aquilegia*-specific without any annotated *A. thaliana* homologs, with most of the remaining genes annotated to be involved in plant defense and basic metabolic functions (Table S4). Since a previous study showed that microRNA2275 (miR2275) precursors and 24-*PHAS* loci are significantly enriched in chromosome 4 compared to other chromosomes in *Aquilegia* (Pokhrel *et al.*, 2021), we also searched for potential microRNA-encoding loci under Q4. MiR2275 is the primary microRNA that triggers phased, secondary, small interfering RNAs (phasiRNAs) of 24 nucleotides in length, and the production of 24-nt phasiRNAs requires both miR2275 copies and 24-*PHAS* loci as their targets (Liu *et al.*, 2020). We found that the region with the highest LOD under Q4 overlaps with the genomic region with the highest density of miR2275 precursors and *24-PHAS* loci, including three of the 11 annotated clusters of miR2275 precursors, and 91 24-*PHAS* loci in total, comprising 31.7% of all 24-*PHAS* loci on chromosome 4 (~45.8 Mb) and 14.1% of such loci in entire genome (~300 Mb) (Pokhrel *et al.*, 2021).

For Q5, there are two genes located within the highest LOD interval that have homologs that are known to function as FM regulators: *AqSEUSS* (*AqSEU*) and *AqARGONAUTE5a* (*AqAGO5a*). In *A. thaliana, SEU* is known to repress *AG* to regulate FM and organ patterning (Pfluger & Zambryski, 2004; Grigorova *et al.*, 2011; Wynn *et al.*, 2014), while in *Aquilegia*, *AqAGO5a* has been identified as a core hub gene associated with early FM development (Min & Kramer, 2020).

Q6 spans 1.5 Mb with 226 genes in total, 170 of which are expressed in the RNAseq dateset (Fig. 4; Table S4). Within the bin of the highest LOD score, there is one gene for which a homolog has been functionally studied: *HANABA TANARU* (*AqHAN*). *HAN* codes for a GATA type zinc finger transcription factor, and in *A. thaliana, HAN* is expressed at the organ boundaries, is known to regulate *WUS* expression, and directly interacts with a number of key genes in FM regulation and primordia initiation (Zhao *et al.*, 2004; Ding *et al.*, 2015). As mentioned above, we have detected significant interaction between Q1 and Q6 (Fig. S5). We found it intriguing that HAN and PTL interact through JAG to control FM morphogenesis in *A. thaliana* (Ding *et al.*, 2015), since their *Aquilegia* homologs are located within the confidence interval of Q1 and Q6, respectively.

Lastly, Q7 also has a very narrow interval of only 2.5 Mb in the genome with 315 genes in total and 242 were expressed in the RNAseq dataset (Table S4). *AqHOMEOBOX GENE 1* (*AqATH1*) is the only gene with a homolog that has been functionally studied that is also located within the highest LOD interval. *AqATH1* is the ortholog of gene *ATH1* in *A. thaliana* (Fig. S8) and belongs to the *BELL1-*like homeodomain gene family. *ATH1* regulates the boundary between the stem and the lateral organs, but is also involved in stem cell regulation in meristems by maintaining the expression of the meristem marker gene *SHOOT MERISTEMLESS* via a self-activation loop (Gómez-Mena & Sablowski, 2008; Li *et al.*, 2012; Cao *et al.*, 2020). *AqATH1* is broadly expressed across the *Aquilegia* FM throughout the early developmental stages (Fig. 5q-t), in all early floral organ primordia (Fig. 5r-t), and at the distal tip of the young lateral organs such as the bracts (Fig. 5q), sepals (Fig. 5r, s), and petals (Fig. 5t).

## DISCUSSION

### *Aquilegia* is an ideal system for studying FM regulation and termination

Over recent decades, we have gained significant insight into various aspects of plant meristem development and function, but the regulation of FMT remains a poorly studied subject. This is despite the fact that FMT is an indispensable process in floral development and variation in FMT timing is a key component of the generation of floral morphological diversity. Progress in understanding the regulation and evolution of FMT is hampered due to the lack of natural variation in floral organ whorl numbers in all of our currently established model systems (and their close relatives), while taxa with such a variation generally lack the genomic and molecular resources to investigate this question further. To this end, *Aquilegia* can be an ideal system for studying FMT because species of *Aquilegia* share relatively low interspecific sequence variation combined with a high degree of interfertility thanks to its recent adaptive radiation (Hodges & Arnold, 1994; Filiault *et al.*, 2018). At the same time, they all share a consistent floral bauplan that only varies in SWN (Munz, 1946), and possess a fully sequenced and well-annotated genome along with RNAi-based methods for functional studies (Kramer, 2009; Filiault *et al.*, 2018). Recognizing that floral SWN is the best available quantitative trait to represent the timing of FMT, we utilized a genetic cross between two sister species differing in SWN, and sought to take the first step to explore the molecular basis of naturally occurring variation in the FMT timing. The mean SWN of *A. brevistyla* and *A. canadensis* is 7.16 and 9.16, respectively, and they do not overlap, while the mean SWN of their F2 progeny was found to encompass the entire range of the parental species (Fig. 2e).

One question we sought to explore was whether the differences in ultimate floral size between the two sister species is reflected by differences in early FM growth dynamics. By analyzing developmental histological series of FMs in the parental species, we detected a subtle yet significant difference in meristem diameter. Although the FMs of both species exhibit an increase in width during their earliest developmental stages and then staying relatively constant during the later stage, the FMs of *A. canadensis* tend to be larger at inception and grow to a larger size, even before differences in SWN are evident (Fig. 3; Table S1). Overall, we observe that 1) the *A. canadensis* FMs are larger in general, 2) have a longer developmental window to increase FM width, and yet, 3) still make five stamens per whorl. There are numerous previous studies showing that an increase in FM diameter is often associated with an increase in floral organ number *per whorl*, rather than an increase in the number of whorls (e.g., Carles *et al.*, 2004; Fan *et al.*, 2014; Chu *et al.*, 2019). Of course, these studies typically rely on mutagenesis or gene over-expression rather than natural variation. This suggests that natural variation in meristem size relies on a greater degree of coordination such that meristem size changes in conjunction with the size of primordia inhibition fields, allowing merosity to stay constant. The current data does not allow us to distinguish between whether the *A. canadensis* FM is growing for a longer period (e.g., perhaps plastochrons are slower, allowing more mass to be accumulated between subsequent whorls) or proliferating at a faster rate. Given what we know about the role of cell division timing in influencing FMT, answering this question is important to understanding the FMT mechanism in *Aquilegia*. Future studies using a recently developed live imaging technique in *Aquilegia* (Min *et al.*, 2021) may allow us to compare growth rates between the initiation of successive whorls in these two species and better characterize this phenomenon.

Another curious observation regarding SWN is that we observed a small portion of individuals in both parental species as well as the F2s that exhibited no variation in SWN, regardless of how many flowers were counted on the plants. In contrast, most other individuals exhibited variation in SWN within an individual plant (Fig. S3). This seems to suggest that there is variation in the robustness of this trait between different individuals. Unfortunately, the fact that there was no significant divergence in this pattern between the parent species meant that we could not map it in the current study, but we hope that examination of within-inflorescence SWN canalization in other *Aquilegia* species will allow the identification of suitable models and the dissection of its genetic basis.

### Variation in the timing of *Aquilegia* FMT is controlled by multiple loci of small effects

We recovered seven major QTL that are responsible for variation in SWN, with one QTL located on each chromosome, and the percent of phenotypic variance explained by each QTL ranging from 3.7% to 8.8% (Fig. 4; Table 1). These results are comparable to previous studies in meristem-related traits of domesticated crops, particularly maize, which also revealed multiple QTL of small effects (Vlăduţu *et al.*, 1999; Upadyayula *et al.*, 2006; Bommert *et al.*, 2013; Thompson *et al.*, 2014, 2015). Interestingly, although all the meristem-related traits measured in maize were highly heritable, the total percentage of variance explained by all the QTL was never higher than 50% (e.g., Bommert *et al.*, 2013; Thompson *et al.*, 2014, 2015), suggesting there are other loci with even smaller effects that were not picked up by the QTL mapping, which is a likely scenario for our current study as well.

We have identified the candidate genes under the QTL (Fig. 4; Table S5), and, further, uncovered novel FM expression patterns of *AqZPR3a* and *AqROXYa*, which were the candidate loci associated with Q2 and Q3, respectively (Fig. 4, 5). In *A. thaliana* and tomato, expression of the *ZPR* genes is restricted to the adaxial region of lateral organs and the central zone of the shoot meristem, and the *ZPR* genes function in both establishing organ polarity and restricting the stem cell domain in the meristems by acting as post-translational suppressors of the class III HD-ZIP abaxial identity genes by inhibiting their homodimerization (Wenkel *et al.*, 2007; Kim *et al.*, 2008; Weits *et al.*, 2019; Xu *et al.*, 2019). However, we have also observed strong expression of *AqZPR3a* in the central epidermal layer of FMs throughout their early developmental stages (Fig. 5f-i), which has not been observed in any previous studies. It will be very interesting to determine whether this expression pattern indicates a novel function or related to known ZPR functions in modulating meristem regulation. In the case of *ROXY* homologs in other models, expression has been found to be restricted to incipient and newly emerged organ primordia (Xing *et al.*, 2005; Li *et al.*, 2009b; Yang *et al.*, 2015), but abaxialized expression such as what was found for *AqROXYa* has not been observed before. In *A. thaliana*, *ROXY1* is known to interact with *PTL* to regulate floral primordium initiation, while in maize, a *ROXY* homolog controls meristem size primordia (Xing *et al.*, 2005; Li *et al.*, 2009b; Yang *et al.*, 2015); either of these functions could be important for controlling FMT in *Aquilegia*.

The Q4 locus is of particular interest because it explains the highest relative percentage of phenotypic variation, but it is also the QTL with the fewest obvious candidate genes to investigate (Table S3, S4). Chromosome 4 of *Aquilegia* appears to have followed a distinct evolutionary path from the rest of the genome and displays many unique features compared to the remaining six chromosomes, including having a higher proportion of genes arrayed in tandem and segmental duplicates, more genetic polymorphism and transposable elements, lower gene density, and reduced gene expression (Filiault *et al.*, 2018; Aköz & Nordborg, 2019). Although the *AqAG1* tandem duplication is included in the 95% Bayesian credible interval, it may be less likely to be the causative gene compared to other genes that were located closer to the highest LOD score marker. The lack of potential candidate genes under Q4 led us to consider other factors besides protein coding genes, leading to the finding that the highest Q4 LOD interval overlaps with the region that harbors the most concentrated density of miR2275 precursors and 24-*PHAS* loci in the entire *Aquilegia* genome (Pokhrel *et al*., 2021). As the primary microRNA that triggers 24-nt phasiRNA, a pathway that is conserved across the angiosperms, miR2275 has been shown to be expressed in the reproductive tissues of various monocot and dicot lineages, particularly in developing anthers (Zhai *et al.*, 2015; Fei *et al.*, 2016; Kakrana *et al.*, 2018; Pokhrel *et al.*, 2020, 2021). However, relatively little is known about 24-nt phasiRNAs in general besides their functions in anthers. Overall, chromosome 4 remains an enigmatic component of the *Aquilegia* genome, so it is intriguing that the QTL is located on this structure. Certainly, it is also possible that the causal gene underlying Q4 is one of the *Aquilegia*-specific loci that did not have a direct *A. thaliana* homolog, which equally applies to the other QTL as well.

The current study is a key first step in identifying a promising list of candidate genes for regulating natural variation in FMT. Next steps in evaluating these candidate genes will include assays of gene function, conducting comparative expression analyses between *A. canadensis* and *A. brevistyla*, and examining sequence variation and patterns of allelic differentiation between populations of these species. Further areas of interest would also include exploring the potential ecological consequences of variation in SWN and FMT between these species and across the genus.

## MATERIALS AND METHODS

### Plant material and growth conditions

*A. brevistyla* and *A. canadensis* seeds were collected from wild populations in Alberta (Canada) and Ithaca (NY, USA), respectively. One *A. canadensis* (pollen recipient) was crossed with one *A. brevistyla* (pollen donor) to generate the F1 generation. Five F1s were self-fertilized to generate the F2 population. All F2 seeds were stratified at 4 °C in the dark for two to four weeks, germinated in wet soil, and transplanted in individual pots. All plants were vernalized at 4°C for two months to induce flowering. The parental and F1 individuals were grown in the greenhouse of the University of California Santa Barbara, and the F2 populations were grown in the greenhouses of Harvard University. All greenhouses used the same light and temperature conditions to achieve a 16h/8h (day/night) photoperiod at 18°C and 13°C.

Seeds of *Aquilegia* x *coerulea* ‘Kiragami’ were purchased from Swallowtail Garden Seeds (Santa Rosa, CA, USA), germinated in wet soil, and grow under the same 18°C /13°C (day/night) condition as described above. Once the plants developed approx. six true leaves, they were transferred into vernalization conditions (16 h daylight at 6 °C and 8 h dark at 6 °C) for three to four weeks, and then moved back to the regular growth conditions to promote flowering.

### Meristem wdith measurement

The entire inflorescences of at least six individuals of each parental species were collected and fixed in FAA (10% formaldehyde, 50% ethanol, 5% acetic acid), and stored at 4°C. Samples were then dehydrated through a graded ethanol series to 100%, transferred to 100% CitriSolv, and embedded in Paraplast Plus (Sigma-Aldrich). Embedded tissues were sectioned to 8 μm thick ribbons with a rotary microtome, stained in 0.1% Toluidine Blue O solution following the protocol described in Ruzin (1990), and mounted in Permount Mounting Medium (Fisher Scientific). Sections were then imaged using the Axio Zoom Microscope at the Harvard Center for Biological Imaging. The width of each floral meristem section was measured using ImageJ. Three to six serial sections were measured for each FM, and at least three FMs were measured for each developmental stage of each species. All FMs that were measured were non-terminal flowers and the number of non-sepal primordia captured by the section was counted.

### Phenotyping

For each plant, the SWN of the terminal flower and lateral flowers 1 to 4 from the first three inflorescences were counted (Fig. 2c). If flowers of these positions in an inflorescence were damaged/undeveloped, flowers at other positions were counted to achieve a total number of five flowers per inflorescence. If an inflorescence produced less than five flowers, all flowers were counted. SWN were counted when the flowers reached approx.1-2 cm in length (Fig. 2d) because at that developmental stage, all the stamens were arranged in vertical rows, which simplified counting.

### Genotyping

Detailed genotyping information can be found in (Edwards *et al.*, 2021). Briefly, the DNA of the two parents that generated the cross was extracted from flash-frozen young leaves using NEBNext Ultra II kit (NEB) and sequenced to ~40x coverage as 150 bp reads on an Illumina MiSeq at the Biological Nanostructures Lab in the California NanoSystem Institute at UC Santa Barbara. DNA of F2s was extracted from silica dried young leaves using Qiagen DNEasy reagents and Magattract beads (Qiagen, Inc.), libraries were prepared following the protocol of RipTide High Throughput Rapid DNA Library Preparation kit (iGenomX, CA, USA). The F2 libraries were pooled and sequenced at the Vincent J. Coates Genomics Sequencing Laboratory (UC Berkeley) using NovaSeq 6000 platform to generate 150bp paired-end reads. Samples were multiplexed to generate about 1-2x coverage. All sequence data are deposited in the Sequence Read Archive under BioProject ID PRJNA720109. Scripts and genotype/phenotype data are available at: https://github.com/anjiballerini/can.x.brev/.

Sequences were aligned to the *A. coerulea* ‘Goldsmith’ v3.1 reference genome (https://phytozome.jgi.doe.gov) using the Burrows-Wheeler aligner (Li & Durbin, 2009) and variable sites in the parents were identified using SAMtools 0.1.19 (Li *et al.*, 2009a) with custom scripts were used to identify the positions and genotypes at which the parents were homozygous for different alleles. These sites were used to assign reads in the F2s as having either *A. canadensis* or *A. brevistyla* ancestry. To determine the genotypes of the F2s, the genome sequences were binned into 0.5 Mb regions with moderate to high recombination frequencies and 1 Mb in regions with low or no recombination, and the frequency of reads with ancestry for each F0 parent was used to determine the genotype of the bin. These bins and genotypes were used as markers to construct a genetic map and conduct QTL mapping. This genotyping method has been implemented in (Filiault *et al.*, 2018; Ballerini *et al.*, 2020; Edwards *et al.*, 2021).

### Mapping

After filtering out individuals and markers with more than 10% of information missing due to sequencing quality, we retained a total of 366 individuals and 620 markers. A genetic map of the seven chromosomes was then constructed following the protocol of the R/qtl package v1.46-2 (Broman *et al.*, 2003), with an error probability rate of 0.001 and “kosambi” map function. Standard interval mapping with Haley-Knott regression (function *scanone*) was used for the initial mapping searching for potential QTL. The best multi-QTL models are produced and selected by using function *stepwiseqtl*, which implement penalties on different interactions and drop one of the current main effects or interactions in each round of model comparison. Interactions among potential QTL and between QTL and covariance were detected with a two-dimensional genome scan (function *scantwo*). Using the estimated positions of QTL from *scanone*, *stepwiseqtl*, and *scantwo* as the input, the positions of QTL were refined by using the function *makeqtl* and *refineqtl*, which then fit with a defined multiple-QTL model (function *fitqtl*) with all detected interactions. F1-parent-of-origin was used as covariance in all the tested mapping models. Position and effect size of QTL were estimated using drop-one-term ANOVA in the best-fitting model. Chromosome diagram with candidate genes (Fig. 4) was produced by using the LinkageMapView (Ouellette *et al.*, 2018).

### *In situ* hybridization of candidate genes

Variable regions of the genes of interest were amplified by PCR (primers in Table S6) from young inflorescence cDNA of *Aquilegia* x *coerulea* ‘Kiragami’. The PCR products were cloned into the pCR™4-TOPO vectors, sequenced to confirm identity, and reverse transcribed using T3 or T7 RNA polymerase and DIG RNA labeling mix (Sigma-Aldrich). Probe qualification and *in situ* hybridization steps followed (Kramer, 2005). Slides were stained in calcofluor white for 5 min before imaging, and pictures were taken using the ZEISS Axio Zoom at the Harvard Center for Biological Imaging.

### Statistical analysis

All statistical analyses (e.g. ANOVA, Tukey’s HSD) were performed using R (version 1.1.456).

### Gene trees

Homologs of *AqROXYa* and *AqATH1* from various taxa were obtained by using BLAST on Phytozome (https://phytozome-next.jgi.doe.gov/). Multiple sequence alignments and neighbor-joining phylogenetic trees were constructed using Geneious Prime (v2021.1.1). We did not construct phylogenetic trees for other candidate genes because their homologs in *A. coerulea* and *A. thaliana* were each other’s reciprocal top BLAST hits.

## AUTHOR CONTRIBUTIONS

YM and EK conceived of and designed the study. EB collected seeds, crossed and sequenced the parental species, and grew and pollinated the F1 individuals. ME and YM grew the F2s. YM conducted all phenotyping. EB, ME, and YM prepared the libraries for sequencing of the F2 individuals and constructed the genetic map. YM did all the data analysis and *in situ* hybridization. EK and SH provided oversight of the study. YM wrote the manuscript with input from the co-authors.

## ACKNOWLEDGEMENT

The authors would like to thank Nicole Bedford, Olivia Meyerson, and Rubén Rellán-Álvarez for discussing QTL mapping analysis; Suresh Pokhrel for providing information on miRNA in *Aquilegia*; Pierre Baduel and Rebecca Povilus for discussing data analysis strategies; the graduate students at the Harvard Statistics Department who volunteered their time for professional and free statistics consultation; Karl Broman for actively answering questions about the R/qtl package on online forums. Funding has been provided by a Simmon’s Award to YM from the Harvard Center for Biological Imaging; a National Science Foundation Graduate Research Fellowship under grant no. DGE1745303 to MBE; and both a NIH Ruth L. Kirschstein National Research Service Award (F32GM103154) and a UC Santa Barbara Harvey Karp Discovery award to ESB. Sequencing was carried out by the DNA Technologies and Expression Analysis Cores at the UC Davis Genome Center, supported by NIH Shared Instrumentation Grant 1S10OD010786-01 and the Biological Nanostructures Lab at UC Santa Barbara.

