## Supplemental figures and tables for "Genetic architecture underlying variation in floral meristem termination in *Aquilegia*"

**Supplementary information:**

Figure S1: SWN and the position of flowers on inflorescences.
Figure S2: F1-parent-of-origin has a significant impact on the distribution of SWN in the respective F2 progeny.
Figure S3: Distribution of the SD of SWN among the parental and F2 populations.
Figure S4: Diagram and summary statistics of the genetic map.
Figure S5: Confirmation of QTL underlying SWN variation.
Figure S6: Effect plots of the markers that have the highest LOD score under each QTL.
Figure S7: Gene phylogeny for *AqROXYa*
Figure S8: Gene phylogeny for *AqATH1*

Table S1. Pairwise comparison of FM widths through early.
Table S2: No significant evidence supporting the presence of a second QTL on any chromosome.
developmental stages.
Table S3: Summary of number of genes under each potential QTL.
Table S4: Expressed genes under Q4.
Table S5: Information on candidate genes.
Table S6: Primers used for constructing *in situ* hybridization probes.

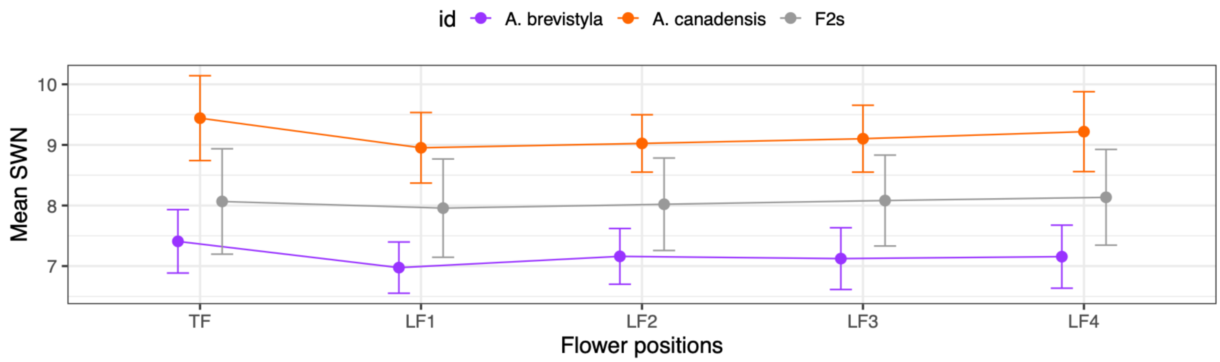


| | | One-way ANOVA | | | | | Tukey's HSD | | | | |
|---|---|---|---|---|---|---|---|---|---|---|---|
| | | Df | Sum Sq | Mean Sq | F value | Pr (>F) | | diff | lwr | upr | p-adj |
| A. brevistyla | flower | 4 | 2.02 | 0.5049 | 4.176 | 0.00325 ** | TF-LF1 | 0.383 | 0.121 | 0.645 | 0.00084 *** |
| | residuals | 130 | 15.72 | 0.1209 | | | | | | | |
| A. canadensis | flower | 4 | 2.387 | 0.5967 | 2.764 | 0.0335 * | TF-LF1 | 0.48 | 0.02 | 0.938 | 0.036 * |
| | residuals | 75 | 16.191 | 0.2159 | | | | | | | |
| F2s | flower | 4 | 1.56 | 0.3909 | 0.875 | 0.479 | | | | | |
| | residuals | 289 | 125.53 | 0.4344 | | | | | | | |

Signif. codes: 0 '***' 0.001 '**' 0.01 '*' 0.05 '.' 0.1 ' ' 1

**Figure S1: SWN and the position of flowers on inflorescences**. Positions of flowers on inflorescences have no significant influence on the SWN among the F2s, but do differ in the parental species, where terminal flowers tend to have higher SWN.

| | Df | Sum Sq | Mean Sq | F value | Pr (>F) |
|---|---|---|---|---|---|
| F1 parent | 4 | 19.40 | 4.850 | 16.32 | 3.39e-12 *** |
| Residuals | 326 | 96.89 | 0.297 | | |

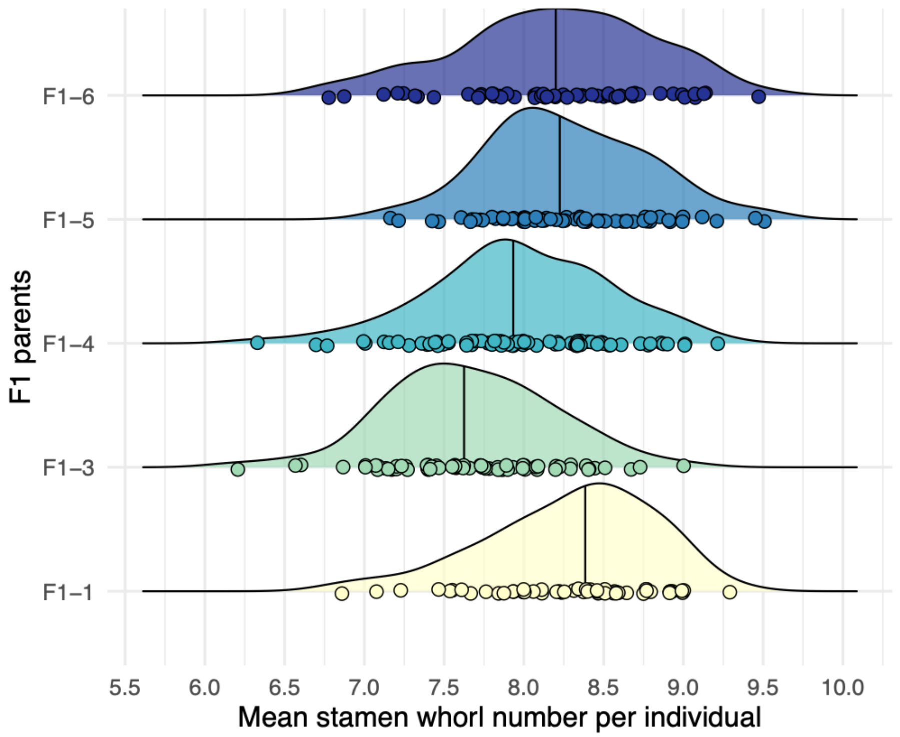


**Figure S2: F1-parent-of-origin has a significant impact on the distribution of SWN in their respective F2 progeny**. One-way ANOVA and density ridgeline plots showing the distribution of SWN of F2s of different F1 parents.

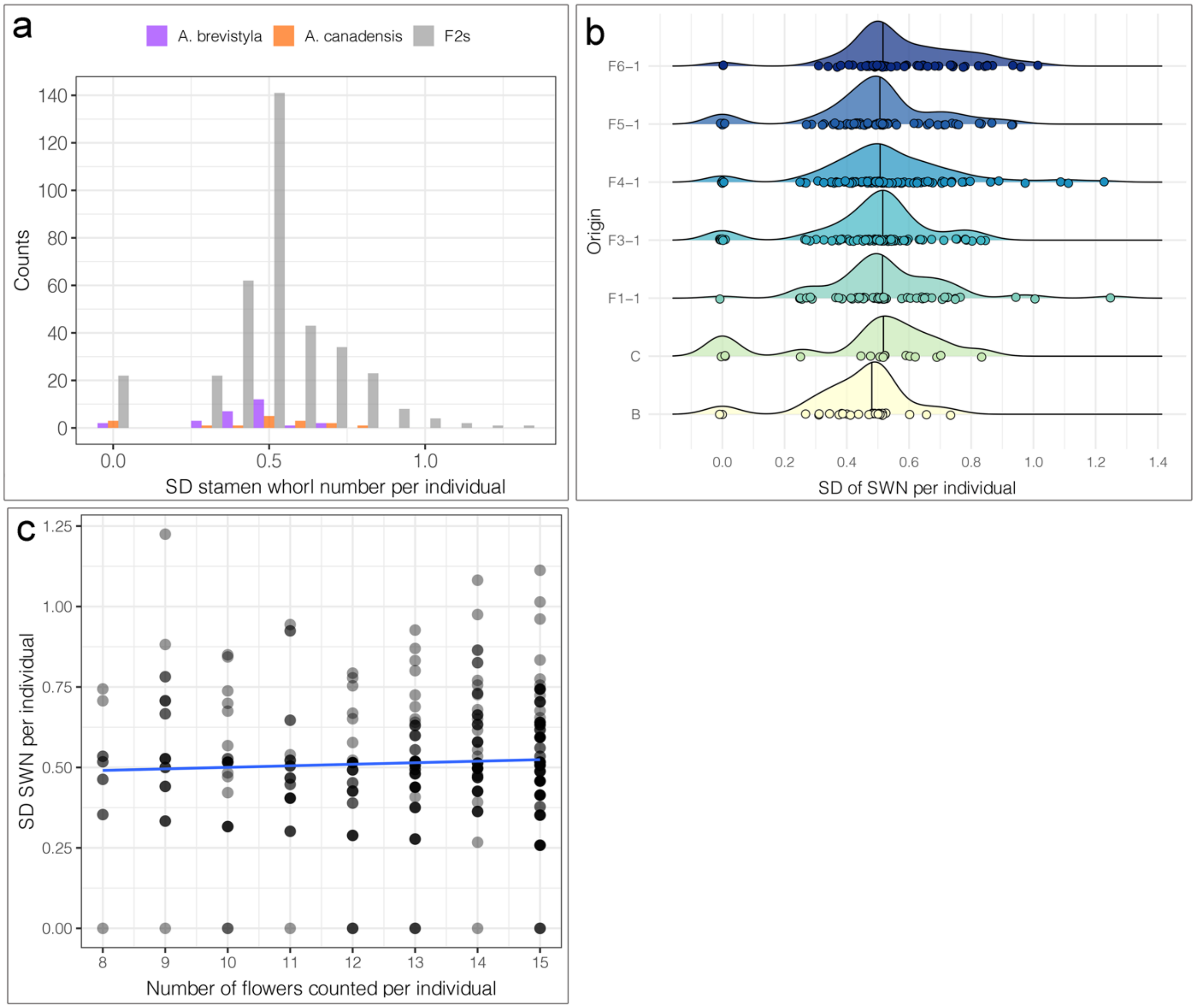


**Figure S3: Distribution of the SD of SWN among the parental and F2 populations.** a. Histogram showing that both parents and the F2s had a small number of individuals showing 0 variation (SD=0) in SWN while the remaining individuals display a large variation in SWN. b. Density ridgeline plots showing the same patterns of SD of SWN distribution regardless of F1-parent-of-origin. c. SD=0 is not an artifact of individuals with fewer flowers counted. There are individuals exhibiting SD=0 in SWN regardless of how many flowers were counted per plant.

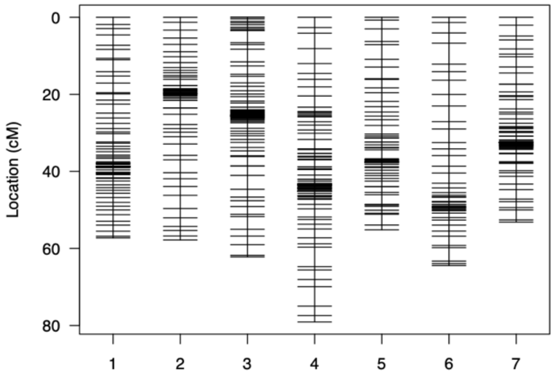


| Linkage group | Number of markers | Length (cM) | Average spacing between markers (cM) | Maximum spacing between markers (cM) |
|---|---|---|---|---|
| 1 | 78 | 55.8 | 0.7 | 3.0 |
| 2 | 66 | 55.4 | 0.9 | 3.3 |
| 3 | 139 | 60.4 | 0.4 | 3.4 |
| 4 | 119 | 76.6 | 0.6 | 4.8 |
| 5 | 71 | 53.3 | 0.8 | 3.7 |
| 6 | 47 | 61.3 | 1.3 | 5.2 |
| 7 | 100 | 51.3 | 0.5 | 2.8 |
| overall | 620 | 414.3 | 0.7 | 5.2 |

**Figure S4: Diagram (left) and summary statistics (right) of the genetic map.**

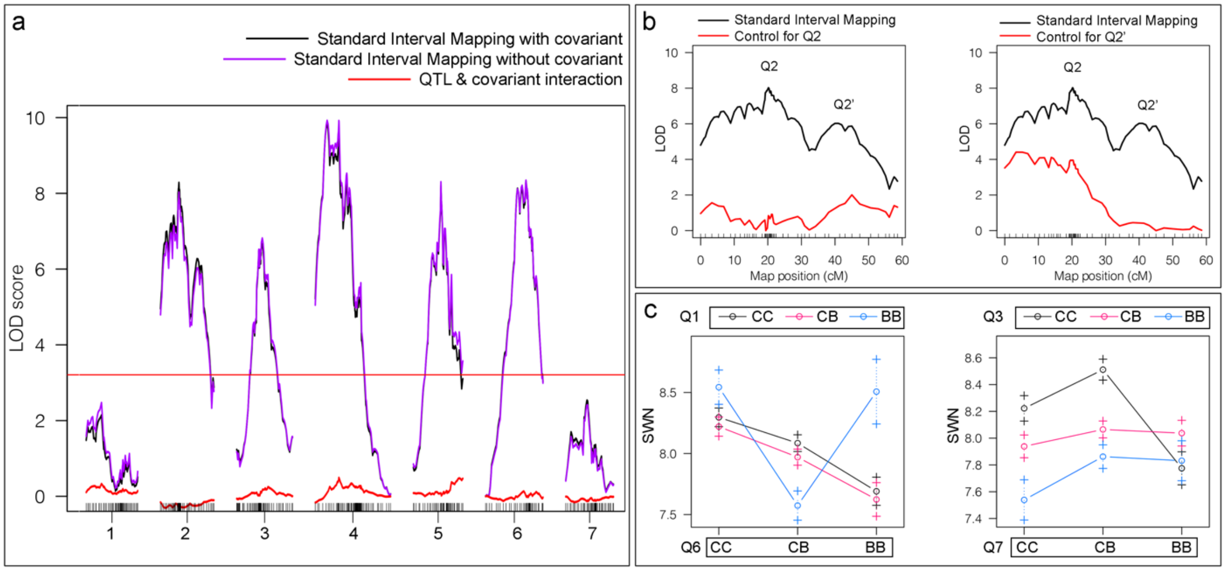


**Figure S5: Confirmation of QTL underlying SWN variation.** (a) No significant interaction between covariant and QTL was detected. (b) Controlling for potential QTL on chromosome 2 did not provide evidence to support the presence of two unlinked QTL. The real QTL on chromosome 2 was named Q2, the potential second QTL was named Q2'. When Q2 was controlled, the evidence for Q2' disappeared, but when Q2' was controlled, the evidence for Q2 stayed significant. (c) QTL interaction between Q1 x Q6, and Q3 x Q7. C: *A. canadensis* allele; B: *A. brevistyla* allele. In (a) and (b), the LOD scores were from the standard interval mapping rather than the full QTL model, which assumes the presence of a single QTL and could not include QTL interactions. This is the reason why the LOD score distributions appear to be a bit different from Fig. 4a.

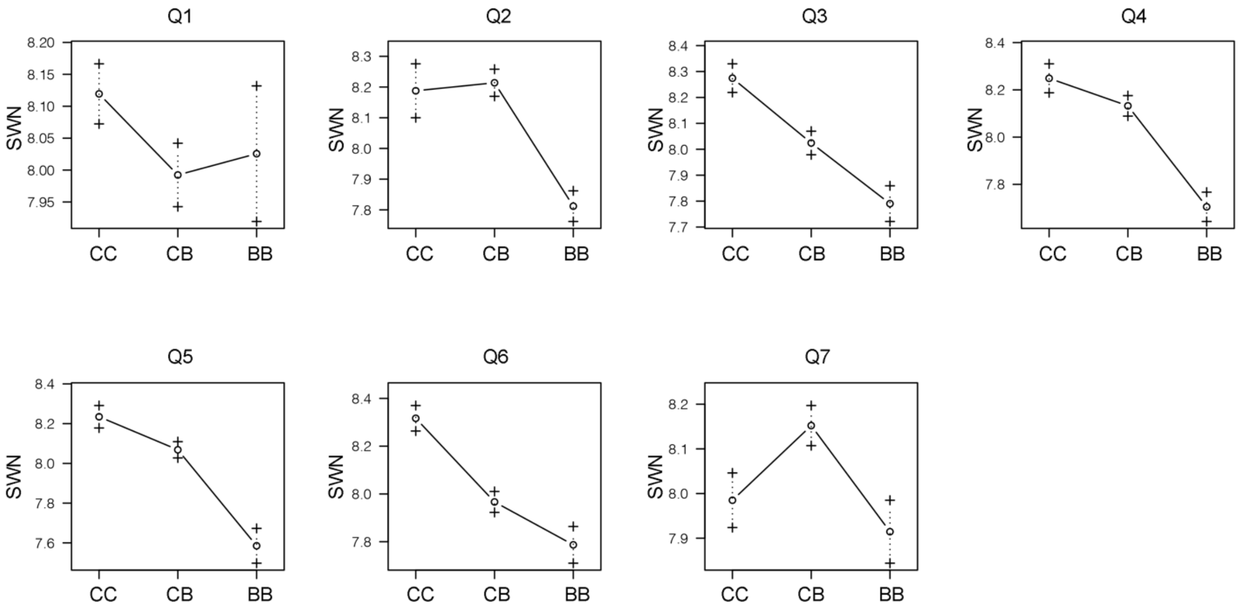


**Figure S6: Effect plots of the markers that have the highest LOD score under each QTL.** C: *A. canadensis* allele; B: *A. brevistyla* allele

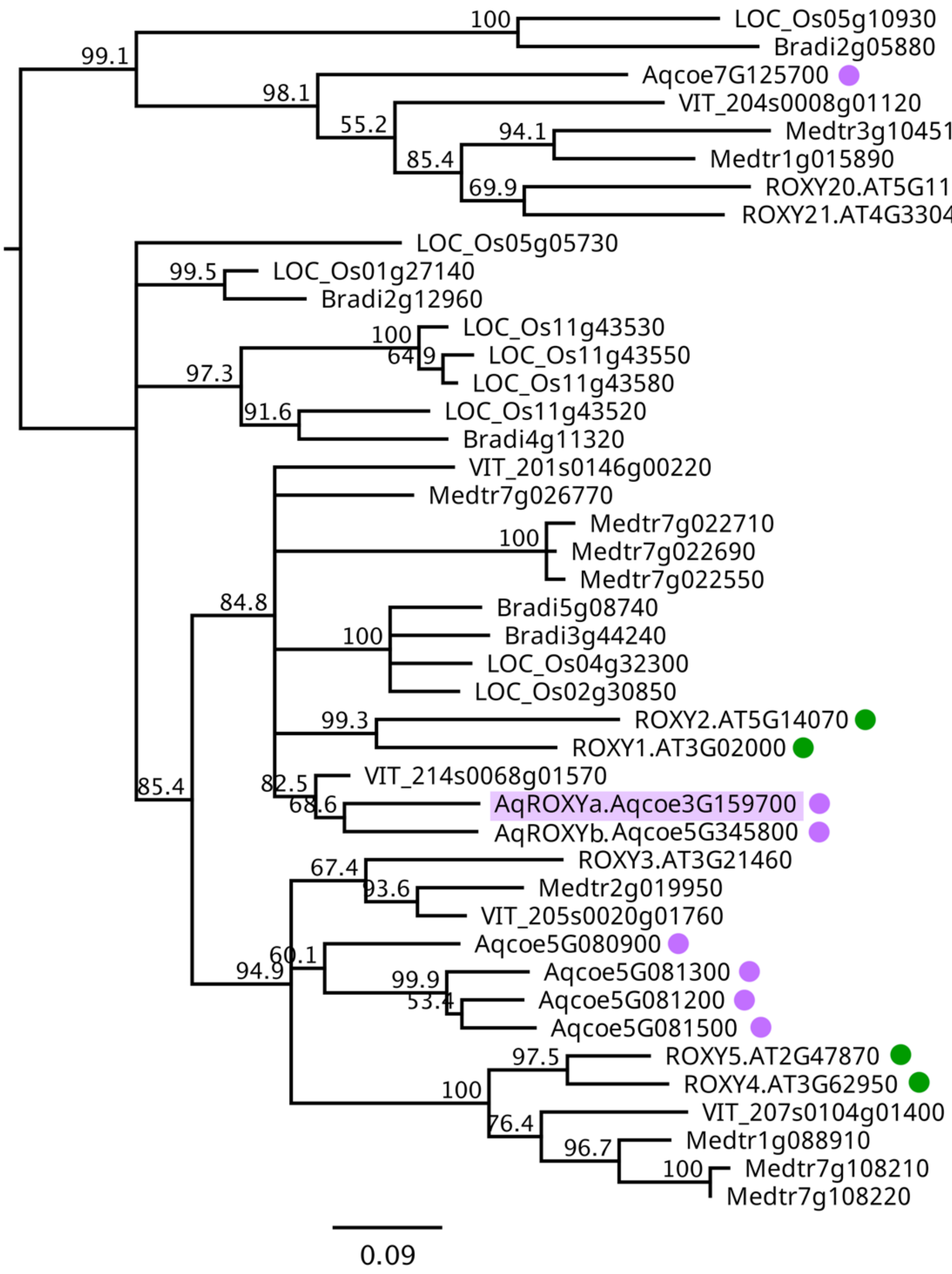


**Figure S7: Gene phylogeny for *AqROXYa***. Neighbor-joining phylogeny of *ROXY* homologs using amino acid alignment. Species included in this phylogeny and the prefix of their gene identifiers: *A. coerulea* ("Aqcoe"), *Vitis vinifera* ("VIT"), *A. thaliana* ("AT"), *Medicago truncatula* ("Medtr"), *Oryza sativa* ("LOC_Os"), and *Brachypodium distachyon* ("Bradi"). All sequences were obtained from Phytozome (https://phytozome-next.jgi.doe.gov/). Homologs in *A. thaliana* and *A. coerulea* are indicated by green and purple dots, respectively. *AqROXYa* is highlighted in purple. This gene phylogeny is consistent with previously published *ROXY* homologs in *A. thaliana* (Li *et al.*, 2009).

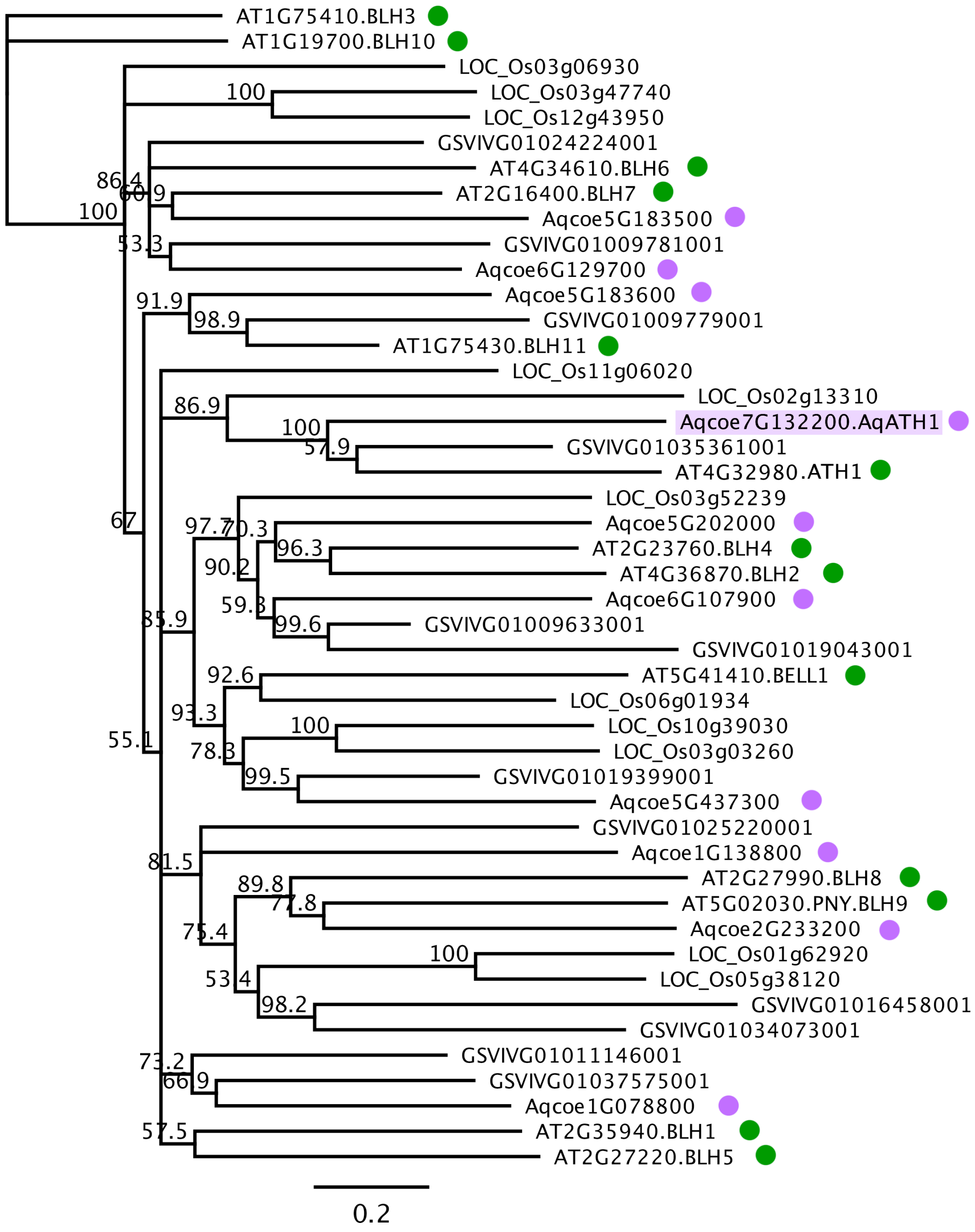


**Figure S8: Gene phylogeny for *AqATH1*.** Neighbor-joining phylogeny of BEL family members. Species included in this phylogeny and the prefix of their gene identifiers: *A. coerulea* ("Aqcoe"), *Vitis vinifera* ("GSVIVG"), *A. thaliana* ("AT"), and *Oryza sativa* ("LOC_Os"). All sequences were obtained from Phytozome (https://phytozome-next.jgi.doe.gov/). Homologs in *A. thaliana* and *A. coerulea* are indicated by green and purple dots, respectively. *AqATH1* is highlighted in purple.

**Table S1. Pairwise comparison of FM widths through early developmental stages.** grp1=group1, grp2=group2 in the pairwise comparison. B: *A. brevistyla*, C: *A. canadensis*. Developmental stages were marked by the number of non-sepal whorls initiated in the FM at a given point, as seen in parenthesis (e.g., 0-2, 3-4, 5-6, 7-8, >8). Comparisons are conducted using the Wilcoxon tests and p-values are adjusted (p.adj) using the Bonferroni test. Pairs that are not significantly different from each other are shaded in light grey. n: number of the sections measured for each group. p.adj.sig: symbols indicating the adjusted p-values in the pairwise comparison; ns: not significant; *: p<0.05; **: p<0.01; ***: p<0.001; ****: p<0.0001.

| | grp1 | grp2 | n (grp1) | n (grp2) | mean (µm, grp1) | mean (µm, grp2) | p | p.adj | p.adj. sig |
|---|---|---|---|---|---|---|---|---|---|
| By stages | B (0~2) | C (0~2) | 37 | 39 | 147.06 | 156.13 | 0.004 | 0.019 | * |
| | B (3~4) | C (3~4) | 20 | 35 | 171.57 | 172.92 | 0.8 | 1 | ns |
| | B (5~6) | C (5~6) | 15 | 15 | 161.60 | 186.78 | 5.16E-08 | 2.58E-07 | **** |
| | B (7~8) | C (7~8) | 16 | 17 | 179.06 | 197.29 | 4.12E-04 | 0.00206 | ** |
| | B (>8) | C (>8) | 20 | 46 | 174.68 | 191.67 | 4.28E-06 | 2.14E-05 | **** |
| By species | B (0~2) | B (3~4) | 37 | 20 | 147.06 | 171.57 | 1.42E-09 | 2.84E-08 | **** |
| | B (0~2) | B (5~6) | 37 | 15 | 147.06 | 161.60 | 3.17E-05 | 6.34E-04 | *** |
| | B (0~2) | B (7~8) | 37 | 16 | 147.06 | 179.06 | 2.54E-08 | 5.08E-07 | **** |
| | B (0~2) | B (>8) | 37 | 20 | 147.06 | 174.68 | 6.46E-10 | 1.29E-08 | **** |
| | B (3~4) | B (5~6) | 20 | 15 | 171.57 | 161.60 | 0.005 | 0.1 | ns |
| | B (3~4) | B (7~8) | 20 | 16 | 171.57 | 179.06 | 0.157 | 1 | ns |
| | B (3~4) | B (>8) | 20 | 20 | 171.57 | 174.68 | 0.425 | 1 | ns |
| | B (5~6) | B (7~8) | 15 | 16 | 161.60 | 179.06 | 8.96E-04 | 0.018 | * |
| | B (5~6) | B (>8) | 15 | 20 | 161.60 | 174.68 | 0.002 | 0.04 | * |
| | B (7~8) | B (>8) | 16 | 20 | 179.06 | 174.68 | 0.381 | 1 | ns |
| | C (0~2) | C (3~4) | 39 | 35 | 156.13 | 172.92 | 1.96E-06 | 3.92E-05 | **** |
| | C (0~2) | C (5~6) | 39 | 15 | 156.13 | 186.78 | 6.93E-12 | 1.39E-10 | **** |
| | C (0~2) | C (7~8) | 39 | 17 | 156.13 | 197.29 | 2.04E-14 | 4.08E-13 | **** |
| | C (0~2) | C (>8) | 39 | 46 | 156.13 | 191.67 | 4.31E-19 | 8.62E-18 | **** |
| | C (3~4) | C (5~6) | 35 | 15 | 172.92 | 186.78 | 2.97E-04 | 5.94E-03 | ** |
| | C (3~4) | C (7~8) | 35 | 17 | 172.92 | 197.29 | 3.12E-09 | 6.24E-08 | **** |
| | C (3~4) | C (>8) | 35 | 46 | 172.92 | 191.67 | 1.56E-08 | 3.12E-07 | **** |
| | C (5~6) | C (7~8) | 15 | 17 | 186.78 | 197.29 | 0.005 | 0.1 | ns |
| | C (5~6) | C (>8) | 15 | 46 | 186.78 | 191.67 | 0.169 | 1 | ns |
| | C (7~8) | C (>8) | 17 | 46 | 197.29 | 191.67 | 0.057 | 1 | ns |

**Table S2: No significant evidence supporting the presence of a second QTL on any chromosome.** The two-dimensional genome scan method scans for all marker pairs of a given chromosome and calculate the LOD scores of the potential two-QTL models based on the marker pairs. The LOD values shown in the table for each chromosome are the highest LOD scores between all pairwise markers of a given chromosome.

*LOD.fv1*: the $\log_{10}$ likelihood ratio comparing the two-QTL full model (including both additive effects and interactions) to the one-QTL model. Large LOD.fv1 scores indicate evidence for a second QTL, allowing for the possibility of interaction.

*LOD.av1*: the $\log_{10}$ likelihood ratio comparing the two-QTL model but only considering additive effects to the one-QTL model. Large LOD.av1 scores indicate evidence for a second QTL, assuming no interaction.

*LOD.int*: the difference between LOD.fv1 and LOD.av1, indicating evidence for interactions between potential QTL.

"10% sig cut off" are the values of LOD.fv1, LOD.av1, and LOD.int above the 90% of their respective distributions generated from 1000 permutations. LOD values that are bigger than the 10% cut off are considered significant.

| | *LOD.fv1* | *LOD.av1* | *LOD.int* |
|---|---|---|---|
| Chromosome 1 | 3.69 | 1.01 | 2.69 |
| Chromosome 2 | 3.86 | 2.31 | 1.55 |
| Chromosome 3 | 2.71 | 2.47 | 0.24 |
| Chromosome 4 | 3.65 | 2.21 | 1.44 |
| Chromosome 5 | 2.83 | 1.71 | 1.12 |
| Chromosome 6 | 3.24 | 1.15 | 2.08 |
| Chromosome 7 | 3.68 | 2.20 | 1.48 |
| 10% sig cut off | 5.99 | 5.04 | 5.25 |

**Table S4: Summary of number of genes under each potential QTL.** Percentages shown in the parentheses are the percentage of the number of genes found in the RNA-seq data (Min & Kramer, 2020) to the total number of genes that are in the respective genomic interval. A gene was considered as expressed if its transcript had more than 1 million read counts in more than 1 samples in the RNA-seq experiment.

| QTL | Chr | size (Mb) | Interval on the genome | No. of genes | Exp at any RNAseq stage |
|---|---|---|---|---|---|
| Q1 | 1 | 6 | 38,000,001..44,000,000 | 964 | 676 (70.12%) |
| Q2 | 2 | 36.5 | 3,500,001.. 40,000,000 | 3242 | 2315 (71.41%) |
| Q3 | 3 | 26.5 | 8,000,001..34,500,000 | 1844 | 1265 (68.60%) |
| Q4 | 4 | 3.8 | 2,000,001..5,800,000 | 383 | 176 (46.21%) |
| Q5 | 5 | 26.5 | 9,000,001..35,500,000 | 1919 | 1393 (72.59%) |
| Q6 | 6 | 1.5 | 6,000,001..7,500,000 | 226 | 170 (75.22%) |
| Q7 | 7 | 2.5 | 7,000,001..9,500,000 | 315 | 242 (76.83%) |

**Table S4 new version: Summary of number of genes, miR2275, and 24-*PHAS* loci under each potential QTL.** Percentages shown in the parentheses are the percentage of the number of genes found in the RNA-seq data (Min & Kramer, 2020) to the total number of genes that are in the respective genomic interval. A gene was considered as expressed if its transcript had more than 1 million read counts in more than 1 samples in the RNA-seq experiment.

| QTL | Chr | size (Mb) | Interval on the genome | No. of genes | Exp at any RNAseq stage | No. of miR2275 precursors (under QTL/on the chr) | No. of 24-PHAS loci (under the QTL/on the chr) |
|---|---|---|---|---|---|---|---|
| Q1 | 1 | 6 | 38,000,001..44,000,000 | 964 | 676 (70.12%) | 0/4 | 10/49 |
| Q2 | 2 | 36.5 | 3,500,001.. 40,000,000 | 3242 | 2315 (71.41%) | 0/0 | 31/56 |
| Q3 | 3 | 26.5 | 8,000,001..34,500,000 | 1844 | 1265 (68.60%) | 3/4 | 28/61 |
| Q4 | 4 | 3.8 | 2,000,001..5,800,000 | 383 | 176 (46.21%) | 4/4 | 91/287 |
| Q5 | 5 | 26.5 | 9,000,001..35,500,000 | 1919 | 1393 (72.59%) | 0/6 | 8/33 |
| Q6 | 6 | 1.5 | 6,000,001..7,500,000 | 226 | 170 (75.22%) | 0/1 | 0/57 |
| Q7 | 7 | 2.5 | 7,000,001..9,500,000 | 315 | 242 (76.83%) | 0/0 | 0/69 |

**Table S5: Expressed genes under Q4.** A gene was considered as expressed if its transcript had more than 1 million read counts in more than 1 samples in the RNA-seq experiment (Min & Kramer, 2020). Best.hit.At: Top BLAST hit A. thaliana gene identifier. Genes under the 1 Mb region that contained the marker with the highest LOD are highlighted in blue.

| Locus ID | Best.hit.At | Symbol | Annotation |
|---|---|---|---|
| Aqcoe4G024000 | AT4G18960 | AG1 | K-box region and MADS-box transcription factor family protein |
| Aqcoe4G024300 | AT3G58770 | AG1 | K-box region and MADS-box transcription factor family protein |
| Aqcoe4G024600 | AT2G42790 | CSY3 | citrate synthase 3 |
| Aqcoe4G024700 | AT5G59840 | | Ras-related small GTP-binding family protein |
| Aqcoe4G024800 | AT5G65750 | | 2-oxoglutarate dehydrogenase, E1 component |
| Aqcoe4G025200 | AT3G54950 | PLA | patatin-like protein 6 |
| Aqcoe4G025400 | AT1G07650 | | Leucine-rich repeat transmembrane protein kinase |
| Aqcoe4G025600 | AT1G29050 | TBL38 | TRICHOME BIREFRINGENCE-LIKE 38 |
| Aqcoe4G026000 | AT3G14840 | LIK1 | Leucine-rich repeat transmembrane protein kinase |
| Aqcoe4G026500 | AT1G53440 | | Leucine-rich repeat transmembrane protein kinase |
| Aqcoe4G026600 | AT1G53440 | | Leucine-rich repeat transmembrane protein kinase |
| Aqcoe4G027000 | AT4G19840 | PP2-A1 | phloem protein 2-A1 |
| Aqcoe4G027100 | AT1G78700 | BEH4 | BES1/BZR1 homolog 4 |
| Aqcoe4G027200 | AT1G53460 | | |
| Aqcoe4G027300 | AT4G18880 | HSFA4A | heat shock transcription factor A4A |
| Aqcoe4G027400 | AT1G78680 | ATGGH2 | gamma-glutamyl hydrolase 2 |
| Aqcoe4G027500 | | | |
| Aqcoe4G027900 | AT1G78680 | ATGGH2 | gamma-glutamyl hydrolase 2 |
| Aqcoe4G028000 | | | |
| Aqcoe4G028600 | AT1G06410 | TPS7 | trehalose-phosphatase/synthase 7 |
| Aqcoe4G028700 | AT1G53380 | | Plant protein of unknown function (DUF641) |
| Aqcoe4G028800 | AT5G61970 | | signal recognition particle-related / SRP-related |
| Aqcoe4G028900 | AT5G41020 | | myb family transcription factor |
| Aqcoe4G029100 | AT2G30950 | FTSH2 | FtsH extracellular protease family |
| Aqcoe4G029200 | AT4G18820 | | AAA-type ATPase family protein |
| Aqcoe4G029300 | AT1G29340 | ATPUB17 | plant U-box 17 |
| Aqcoe4G029700 | | | |
| Aqcoe4G030000 | AT2G42770 | | Peroxisomal membrane 22 kDa (Mpv17/PMP22) family protein |
| Aqcoe4G031000 | AT3G19720 | ARC5 | P-loop containing nucleoside triphosphate hydrolases superfamily protein |
| Aqcoe4G031200 | | | |
| Aqcoe4G031300 | AT4G18800 | RABA1D | RAB GTPase homolog A1D |
| Aqcoe4G031600 | AT1G11330 | | S-locus lectin protein kinase family protein |
| Aqcoe4G031700 | AT2G42760 | | |
| Aqcoe4G031800 | | | |
| Aqcoe4G031900 | AT2G42750 | | DNAJ heat shock N-terminal domain-containing protein |
| Aqcoe4G032000 | | | |
| Aqcoe4G032200 | AT5G45760 | | Transducin/WD40 repeat-like superfamily protein |
| Aqcoe4G032300 | AT4G18780 | CESA8 | cellulose synthase family protein |
| Aqcoe4G032400 | AT3G14067 | | Subtilase family protein |
| Aqcoe4G032600 | AT5G01750 | | Protein of unknown function (DUF567) |
| Aqcoe4G032900 | AT2G27260 | | Late embryogenesis abundant (LEA) hydroxyproline-rich glycoprotein family |
| Aqcoe4G033500 | AT5G13690 | NAGLU | alpha-N-acetylglucosaminidase family / NAGLU family |
| Aqcoe4G034000 | AT5G22870 | | Late embryogenesis abundant (LEA) hydroxyproline-rich glycoprotein family |
| Aqcoe4G034100 | AT4G35490 | MRPL11 | mitochondrial ribosomal protein L11 |

| | | | |
|---|---|---|---|
| Aqcoe4G034500 | AT4G18750 | DOT4 | Pentatricopeptide repeat (PPR) superfamily protein |
| Aqcoe4G034700 | | | |
| Aqcoe4G034900 | AT4G18750 | DOT4 | Pentatricopeptide repeat (PPR) superfamily protein |
| Aqcoe4G035000 | AT5G35370 | | S-locus lectin protein kinase family protein |
| Aqcoe4G035500 | AT5G43470 | HRT | Disease resistance protein (CC-NBS-LRR class) family |
| Aqcoe4G035600 | AT1G78770 | APC6 | anaphase promoting complex 6 |
| Aqcoe4G035700 | AT4G14440 | ATECI3 | 3-hydroxyacyl-CoA dehydratase 1 |
| Aqcoe4G035800 | AT3G14570 | ATGSL04 | glucan synthase-like 4 |
| Aqcoe4G035900 | AT3G25780 | AOC3 | allene oxide cyclase 3 |
| Aqcoe4G036000 | AT2G30970 | ASP1 | aspartate aminotransferase 1 |
| Aqcoe4G036100 | AT5G16860 | | Tetratricopeptide repeat (TPR)-like superfamily protein |
| Aqcoe4G036200 | AT3G59500 | | Integral membrane HRF1 family protein |
| Aqcoe4G036300 | AT5G45780 | | Leucine-rich repeat protein kinase family protein |
| Aqcoe4G036400 | AT2G30980 | BIL1 | SHAGGY-related protein kinase dZeta |
| Aqcoe4G036600 | AT3G58690 | | Protein kinase superfamily protein |
| Aqcoe4G036700 | AT5G26830 | | Threonyl-tRNA synthetase |
| Aqcoe4G036800 | AT4G20020 | | |
| Aqcoe4G036900 | | | |
| Aqcoe4G038100 | | | |
| Aqcoe4G038200 | | | |
| Aqcoe4G038400 | AT1G06620 | | 2-oxoglutarate (2OG) and Fe(II)-dependent oxygenase superfamily protein |
| Aqcoe4G038600 | AT2G34960 | CAT5 | cationic amino acid transporter 5 |
| Aqcoe4G039000 | AT4G17760 | | damaged DNA binding;exodeoxyribonuclease IIIs |
| Aqcoe4G039600 | AT5G48620 | | Disease resistance protein (CC-NBS-LRR class) family |
| Aqcoe4G040000 | AT3G06240 | | F-box family protein |
| Aqcoe4G040200 | AT5G48620 | | Disease resistance protein (CC-NBS-LRR class) family |
| Aqcoe4G040300 | AT2G42690 | | alpha/beta-Hydrolases superfamily protein |
| Aqcoe4G040400 | AT1G53530 | | Peptidase S24/S26A/S26B/S26C family protein |
| Aqcoe4G040500 | AT5G45840 | | Leucine-rich repeat protein kinase family protein |
| Aqcoe4G040600 | AT1G78810 | | |
| Aqcoe4G040700 | AT2G42670 | | Protein of unknown function (DUF1637) |
| Aqcoe4G040800 | | | |
| Aqcoe4G040900 | AT3G14910 | | |
| Aqcoe4G041300 | AT5G17540 | | HXXXD-type acyl-transferase family protein |
| Aqcoe4G042300 | AT5G17540 | | HXXXD-type acyl-transferase family protein |
| Aqcoe4G042400 | AT1G09850 | XBCP3 | xylem bark cysteine peptidase 3 |
| Aqcoe4G042500 | | | |
| Aqcoe4G042700 | AT3G58640 | | Mitogen activated protein kinase kinase kinase-related |
| Aqcoe4G043000 | AT1G14990 | | |
| Aqcoe4G043100 | AT1G29195 | | |
| Aqcoe4G043200 | AT5G54750 | | Transport protein particle (TRAPP) component |
| Aqcoe4G043300 | AT3G04600 | | Nucleotidylyl transferase superfamily protein |
| Aqcoe4G043400 | AT3G04040 | | |
| Aqcoe4G043700 | AT3G58630 | | sequence-specific DNA binding transcription factors |
| Aqcoe4G043800 | AT2G42610 | LSH10 | Protein of unknown function (DUF640) |
| Aqcoe4G043900 | AT1G17020 | ATSRG1 | senescence-related gene 1 |
| Aqcoe4G044000 | | | |
| Aqcoe4G044200 | AT1G29170 | SCAR3 | Encodes a member of the SCAR family. |
| Aqcoe4G045300 | AT5G32440 | | Ubiquitin system component Cue protein |
| Aqcoe4G045600 | AT1G06630 | | F-box/RNI-like superfamily protein |
| Aqcoe4G045700 | AT5G10770 | | Eukaryotic aspartyl protease family protein |
| Aqcoe4G046200 | AT5G10870 | ATCM2 | chorismate mutase 2 |
| Aqcoe4G046500 | AT3G14940 | ATPPC3 | phosphoenolpyruvate carboxylase 3 |

| | | | |
|---|---|---|---|
| Aqcoe4G047100 | AT1G29930 | AB140 | chlorophyll A/B binding protein 1 |
| Aqcoe4G047300 | | | |
| Aqcoe4G047400 | AT4G10780 | | LRR and NB-ARC domains-containing disease resistance protein |
| Aqcoe4G047500 | AT1G29930 | AB140 | chlorophyll A/B binding protein 1 |
| Aqcoe4G047600 | AT1G29120 | | Hydrolase-like protein family |
| Aqcoe4G047700 | | | |
| Aqcoe4G047800 | | | |
| Aqcoe4G048000 | AT1G16890 | UBC13B | ubiquitin-conjugating enzyme 36 |
| Aqcoe4G048200 | AT2G42580 | TTL3 | tetratricopetide-repeat thioredoxin-like 3 |
| Aqcoe4G048400 | AT1G22870 | | Protein kinase family protein with ARM repeat domain |
| Aqcoe4G048700 | AT5G61190 | | putative endonuclease or glycosyl hydrolase with C2H2-type zinc finger domain |
| Aqcoe4G048800 | AT4G16130 | ARA1 | arabinose kinase |
| Aqcoe4G049500 | AT2G15480 | UGT73B5 | UDP-glucosyl transferase 73B5 |
| Aqcoe4G049700 | AT4G14850 | LOI1 | Pentatricopeptide repeat (PPR) superfamily protein |
| Aqcoe4G049800 | AT4G17410 | | DWNN domain, a CCHC-type zinc finger |
| Aqcoe4G050000 | AT4G33355 | | Bifunctional inhibitor/lipid-transfer protein/seed storage 2S albumin superfamily protein |
| Aqcoe4G050300 | AT1G61180 | | LRR and NB-ARC domains-containing disease resistance protein |
| Aqcoe4G050700 | AT3G09270 | GSTU8 | glutathione S-transferase TAU 8 |
| Aqcoe4G050800 | AT3G58610 | | ketol-acid reductoisomerase |
| Aqcoe4G050900 | AT3G14980 | ROS | Acyl-CoA N-acyltransferase with RING/FYVE/PHD-type zinc finger protein |
| Aqcoe4G051100 | AT5G45900 | APG7, | ThiF family protein |
| Aqcoe4G051300 | AT1G53250 | | |
| Aqcoe4G051400 | | | |
| Aqcoe4G051500 | | | |
| Aqcoe4G051600 | | | |
| Aqcoe4G051700 | AT1G78750 | | F-box/RNI-like superfamily protein |
| Aqcoe4G051900 | AT3G59200 | | F-box/RNI-like superfamily protein |
| Aqcoe4G052100 | AT2G31130 | | |
| Aqcoe4G052600 | AT4G19050 | | NB-ARC domain-containing disease resistance protein |
| Aqcoe4G052900 | AT3G50120 | | Plant protein of unknown function (DUF247) |
| Aqcoe4G053000 | AT2G34090 | MEE18 | maternal effect embryo arrest 18 |
| Aqcoe4G053100 | AT3G59010 | PME61 | pectin methylesterase 61 |
| Aqcoe4G053200 | AT1G56000 | | FAD/NAD(P)-binding oxidoreductase family protein |
| Aqcoe4G053300 | AT1G56000 | | FAD/NAD(P)-binding oxidoreductase family protein |
| Aqcoe4G053600 | AT5G44430 | PDF1.2c | plant defensin 1.2C |
| Aqcoe4G053900 | AT1G56000 | | FAD/NAD(P)-binding oxidoreductase family protein |
| Aqcoe4G054200 | AT5G44430 | PDF1.2c | plant defensin 1.2C |
| Aqcoe4G054300 | AT2G47730 | ATGSTF8 | glutathione S-transferase phi 8 |
| Aqcoe4G054400 | AT1G55980 | | FAD/NAD(P)-binding oxidoreductase family protein |
| Aqcoe4G054700 | | | |
| Aqcoe4G054800 | AT1G17120 | CAT8 | cationic amino acid transporter 8 |
| Aqcoe4G054900 | AT5G48620 | | Disease resistance protein (CC-NBS-LRR class) family |
| Aqcoe4G055300 | AT5G44430 | PDF1.2c | plant defensin 1.2C |
| Aqcoe4G055400 | | | |
| Aqcoe4G055600 | AT5G44430 | PDF1.2c | plant defensin 1.2C |
| Aqcoe4G055800 | AT5G44430 | PDF1.2c | plant defensin 1.2C |
| Aqcoe4G055900 | AT5G44430 | PDF1.2c | plant defensin 1.2C |
| Aqcoe4G056100 | AT3G25750 | | F-box family protein with a domain of unknown function (DUF295) |

| | | | |
|---|---|---|---|
| Aqcoe4G056200 | AT5G44430 | PDF1.2c | plant defensin 1.2C |
| Aqcoe4G056300 | AT5G44430 | PDF1.2c | plant defensin 1.2C |
| Aqcoe4G056500 | AT1G16670 | | Protein kinase superfamily protein |
| Aqcoe4G056800 | | | |
| Aqcoe4G056900 | AT1G29050 | TBL38 | TRICHOME BIREFRINGENCE-LIKE 38 |
| Aqcoe4G057100 | AT3G25690 | CHUP1 | Hydroxyproline-rich glycoprotein family protein |
| Aqcoe4G057300 | AT1G78830 | | Curculin-like (mannose-binding) lectin family protein |
| Aqcoe4G057400 | AT1G78850 | | D-mannose binding lectin protein with Apple-like carbohydrate-binding domain |
| Aqcoe4G057900 | AT5G51040 | | |
| Aqcoe4G058000 | AT3G23760 | | |
| Aqcoe4G058100 | | | |
| Aqcoe4G058300 | AT5G20040 | ATIPT9 | isopentenyltransferase 9 |
| Aqcoe4G058400 | AT5G45910 | | GDSL-like Lipase/Acylhydrolase superfamily protein |
| Aqcoe4G058500 | AT1G53240 | mMDH1 | Lactate/malate dehydrogenase family protein |
| Aqcoe4G058600 | AT1G06230 | GTE4 | global transcription factor group E4 |
| Aqcoe4G058700 | AT1G50410 | FRG | SNF2 domain-containing protein / helicase domain-containing protein /zinc finger protein-related |
| Aqcoe4G058900 | AT5G44430 | PDF1.2c | plant defensin 1.2C |
| Aqcoe4G059300 | AT1G73170 | | P-loop containing nucleoside triphosphate hydrolases superfamily protein |
| Aqcoe4G059500 | AT1G73170 | | P-loop containing nucleoside triphosphate hydrolases superfamily protein |
| Aqcoe4G059800 | AT4G18480 | CH-42 | P-loop containing nucleoside triphosphate hydrolases superfamily protein |
| Aqcoe4G059900 | AT2G33980 | NUDT22 | nudix hydrolase homolog 22 |
| Aqcoe4G060500 | AT3G24800 | PRT1 | proteolysis 1 |
| Aqcoe4G060600 | | | |
| Aqcoe4G060700 | | | |
| Aqcoe4G060900 | | | |
| Aqcoe4G061000 | | | |
| Aqcoe4G061200 | AT1G53210 | | sodium/calcium exchanger family protein / calcium-binding EF hand family protein |
| Aqcoe4G061400 | AT5G42820 | U2AF35B | Zinc finger C-x8-C-x5-C-x3-H type family protein |
| Aqcoe4G061800 | AT5G15380 | DRM1 | domains rearranged methylase 1 |
| Aqcoe4G062100 | AT1G11950 | | Transcription factor jumonji (jmjC) domain-containing protein |

**Table S6: Candidate genes under QTL**.

| Chr | Locus ID | Gene name | Genomic location (5’) |
|---|---|---|---|
| 1 | Aqcoe1G411100 | *AqLAS* | 39,117,500 |
| 1 | Aqcoe1G456700 | *AqPTL* | 42,021,250 |
| 2 | Aqcoe2G139400 | *AqZPR3a* | 12,741,500 |
| 3 | Aqcoe3G159700 | *AqROXYa* | 13,927,500 |
| 4 | Aqcoe4G024000 | *AqAG1* | 2,002,500 |
| 5 | Aqcoe5G235700 | *AqSEU* | 15,620,000 |
| 5 | Aqcoe5G237200 | *AqAGO5a* | 15,842,500 |
| 6 | Aqcoe6G121000 | *AqHAN* | 6,619,000 |
| 7 | Aqcoe7G132200 | *AqATH1* | 8,222,500 |

**Table S7: Primers used for constructing in situ hybridization probes.**

| Locus ID | Gene name | Forward primer (5’to 3’) | Reverse primer (5’ to 3’) | Product size |
|---|---|---|---|---|
| Aqcoe2G057900 | *AqWUS* | TGTCGAGCCATATCCATTTTTCAAC | TCATGCATGATGTTATCAGTCCTTTG | 300 bp |
| Aqcoe2G139400 | *AqZPR3a* | CAGAGCTTTACTTGAGGAATTTG | GAACTCTTGGGATTTGGAGA | 196 bp |
| Aqcoe3G159700 | *AqROXYa* | AAATACCAAACACACCAACT | TTGACATGTATGACAACTCC | 220 bp |
| Aqcoe7G132200 | *AqATH1* | GGCAGTTCTAGTTCTATTGC | TGAAGCTGTTGAAACATTTA | 147 bp |
